# Frequency-Specific Operant Learning in Neurofeedback Reveals Distinct Cortical Mechanisms: Evidence from Double-Blind ERSP and ERP Dissociations

**DOI:** 10.64898/2026.04.13.718260

**Authors:** Andrew Hill

**Affiliations:** Peak Brain Institute, Los Angeles, CA

## Abstract

**Background:** Neurofeedback reliably alters EEG activity, but two questions remain unresolved: what cortical mechanism underlies reward-contingent learning, and whether training produces durable resting-state change rather than transient within-session shifts. No study has examined reward-locked event-related spectral perturbations (ERSP) under double-blind, active-placebo-controlled conditions.

**Methods:** Forty participants underwent five training sessions of single-channel EEG biofeedback (C3 SMR 12–15 Hz, *n* = 8; C3 Beta 15–18 Hz, *n* = 8; C4 SMR 12–15 Hz, *n* = 8; active-placebo sham, *n* = 16), plus a retention session a median of 29 days post-training, with concurrent 64-channel EEG recording. ERSP was computed from reward-locked epochs (approximately 600–700 trials per session) using Morlet wavelets (3–40 Hz) across four sessions. Resting-state alpha trajectories were modeled with growth-curve mixed-effects models on the pipeline’s fixed 8–12 Hz alpha band, with an IAF-anchored reanalysis (IAF ± 2 Hz) as a robustness check.

**Results:** Under double-blind active-placebo control, SMR (but not Beta) neurofeedback produced a durable elevation of resting eyes-closed alpha persisting to the one-month follow-up (median 29 days). An LME growth curve confirmed SMR-specific accumulation (C3 SMR × Session *β* = 1.44, *p* = 0.004; C4 SMR *β* = 1.24, *p* = 0.012; Beta and sham flat), robust to IAF-anchored reanalysis (C3 SMR *β* = 1.21, *p* = 0.006; C4 SMR *β* = 1.10, *p* = 0.013) and present in 13 of 16 (81%) SMR participants versus 7 of 16 sham. Critically, the Beta group showed the largest immediate reward-locked ERD (*d* = −2.38 vs sham) yet the weakest consolidation, dissociating acute control from durable plasticity.

Active groups produced frequency-specific event-related desynchronization (ERD) in the rewarded band (pooled Active vs Sham *d* = −1.23, *p*_adj_ = 0.001; C3 Beta and C4 SMR FDR-significant, |*d*| ≥ 1.12; C3 SMR trended, *d* = −0.80, *p*_adj_ = 0.081), absent in sham. A double dissociation emerged at C3: beta training produced the strongest ERD (*d* = −2.38), whereas C3 SMR training produced the largest P2 suppression (*d* = −1.33, BF_01_ = 0.10; smaller P2 at the trained electrode), consistent with distinct frequency-specific operant signatures. ERD magnitude did not track within-session resting shifts (*r* = −0.09, *p* = 0.67) and showed no detectable association with durable change at the individual level, consistent with consolidation as a property distinct from acute control. An ICA-based sensitivity analysis confirmed convergence of all primary findings.

**Conclusions:** Neurofeedback engages frequency-specific, contingency-dependent cortical mechanisms whose consolidation profiles differ by protocol. The Beta arm controls the rhythm most strongly during training yet does not consolidate; both SMR arms show weaker acute control but durable resting-state growth. Consolidability, not session count or acute control, is the property that distinguishes protocols. The ERD–P2 dissociation suggests that beta and SMR training engage distinct frequency-specific mechanisms (C3 Beta: stronger spectral ERD with preserved P2; C3 SMR: moderate ERD with suppressed P2 at the trained site) with different capacities for offline consolidation. These findings support a multi-timescale model in which durable plasticity emerges only when the engaged circuit supports offline consolidation, a property not predicted by the magnitude of acute reward-locked control.

## 1. Introduction

Neurofeedback (the provision of real-time feedback about one’s own brain activity) has demonstrated moderate-to-large effect sizes in some meta-analyses (Arns et al., 2009; Van Doren et al., 2019), though well-controlled trials with probably-blinded outcomes have produced smaller effects in ADHD (Cortese et al., 2016). Yet the field remains uncertain about *how* the brain integrates a reward signal yoked to its own neural oscillations and produces lasting changes in cortical physiology. Large sham-controlled trials have produced equivocal efficacy results (Schönenberg et al., 2017; Arnold et al., 2021), in part because the field lacks a mechanistic account to guide protocol design. Sitaram et al. (2017) identified this gap as a central priority; Thibault and Raz (2017) argued that many reported effects could be entirely attributable to placebo. While the field debates whether neurofeedback outperforms sham on behavioral outcomes, no study has examined what occurs in the brain at the moment of reward delivery during training itself.

A separate, underappreciated gap concerns the distinction between within-session control and durable change. The dominant outcome metric in neurofeedback research is whether a participant can move the target band during training. But such control is a state, not a trait: for neurofeedback to produce lasting clinical benefit, training must durably alter the resting brain after feedback stops. Most neurofeedback studies lack both the active-placebo control needed to rule out non-specific resting-state effects and the post-training follow-up needed to distinguish consolidation from transient practice. The clinically relevant question is therefore not only *how* the brain responds to reward-contingent feedback during training, but whether and under what conditions those responses consolidate into lasting changes in resting-state physiology.

The operant conditioning framework provides a testable hypothesis: if the brain learns to control its own oscillatory power through reward-contingent feedback, then measurable, frequency-specific cortical responses should emerge, time-locked to reward delivery. Sterman (1977, 2000) demonstrated that cats and humans could be operantly conditioned to increase sensorimotor rhythm (SMR, 12–15 Hz) power when rewarded for doing so. Fetz (1969, 2007) established that individual cortical neurons could be conditioned through contingent reward. Ros et al. (2010) have argued that neurofeedback engages the same neuroplasticity mechanisms underlying motor learning, and Sherlin et al. (2011; updated by Kerson, Sherlin, & Davelaar, 2025) extended this reasoning to oscillatory control. Davelaar (2018) formalized the first two stages of a multi-stage computational theory (implicit operant learning and structural thalamocortical change); a subsequent synthesis extends this with a third stage of interoceptive homeostasis (Kerson, Sherlin & Davelaar, 2025). Yet until recently, the analytical tools available have been poorly suited to visualizing frequency-specific learning signals with sufficient temporal and spectral resolution.

Event-Related Spectral Perturbation (ERSP) preserves both time and frequency simultaneously, revealing how oscillatory power evolves on millisecond timescales around significant events (Makeig, 1993; Pfurtscheller & Lopes da Silva, 1999; Grandchamp & Delorme, 2011). The distinction between phase-locked and non-phase-locked responses (Tallon-Baudry et al., 1996) is central to this framework. The ERSP framework describes event-related desynchronization (ERD) and synchronization (ERS), transient decreases or increases in band-limited oscillatory power, providing the resolution needed to ask: *Does the brain produce a frequency-specific, temporally-precise oscillatory response contingent on the learning signal, only when feedback is genuinely yoked to the target neural activity?*

Answering this question rigorously requires overcoming a substantial methodological hurdle: blinding. Unlike pharmacological trials, where inert placebos are standard, blinding neurofeedback participants to whether they are receiving real or sham feedback is notoriously difficult. Schönenberg et al. (2017) and Ros et al. (2020) have found that most “sham” conditions fail rigorous scrutiny: they either provide intermittent real feedback, lack temporal coherence, or employ crude pseudorandom sequences that participants can distinguish. Thibault et al. (2016) showed that blinding failure could entirely account for reported effects. Arnold et al. (2013, 2021) and Sorger et al. (2019) have articulated standards for active-placebo control: the sham must provide real-time, artifact-merged EEG data with matched temporal resolution and amplitude scaling, with feedback contingency hidden, such that participants cannot reliably distinguish active from sham training. Only with such controls in place can one credibly claim that observed changes in brain activity are contingent on the *veridical* learning signal.

The present study addresses these gaps by examining both the brain’s *automatic*, reward-locked spectral response during operant conditioning and the trajectory of resting-state change across sessions and into a multi-week follow-up, without instructing subjects to perform any specific mental task. Unlike prior work that examines ERPs or spectral changes *before and after* neurofeedback, including recent double-blind, placebo-controlled ERP studies of sensorimotor training (Dousset et al., 2024, 2025) and earlier frequency-specific designs (Egner & Gruzelier, 2001, 2004), that studies ERD during *voluntary* motor imagery aided by feedback (Muraoka et al., 2024), or that evaluates frequency specificity using only tonic power from a single training frequency (Dessy et al., 2020), we examine the reward-locked ERSP itself. Four groups (N = 40: C3 SMR n=8, C3 Beta n=8, C4 SMR n=8, Sham n=16) underwent double-blind, active-placebo-controlled neurofeedback with concurrent 64-channel EEG across five training sessions plus a retention visit (median 29 days post-training). We hypothesized that active groups would produce frequency-specific ERD in the rewarded band, time-locked to reward delivery, absent in sham. We additionally tracked resting-state alpha trajectories across sessions and into the follow-up to assess whether protocol-specific acute effects consolidate into durable resting-state change. These data were originally analyzed descriptively (Hill, 2012); we reanalyze them here using cluster-based permutation testing, Bayesian inference, mixed-effects models, and standardized effect sizes.

## 2. Methods

### 2.1 Participants and Design

Forty healthy adults with no prior neurofeedback experience were recruited from the University of California, Los Angeles undergraduate community. Participants were randomly assigned to one of four groups by an unaffiliated researcher: C3 SMR (n = 8; reward 12–15 Hz at C3), C3 Beta (n = 8; reward 15–18 Hz at C3), C4 SMR (n = 8; reward 12–15 Hz at C4), or active-placebo sham (n = 16). All 40 participants were included in analyses. Data were collected as part of the first author’s doctoral research (Hill, 2012); the present ERSP-focused reanalysis was designed in 2025– 2026 using statistical methods not available at the time of original data collection, with the analysis plan documented prior to re-accessing raw BDF files. Because the first author’s 2012 dissertation analyzed these data descriptively, the present reanalysis is not a pre-registered replication. The pre-documented analysis plan mitigates but does not eliminate prior-exposure bias; a prospective replication is the natural next step. The study was approved by the UCLA Institutional Review Board; all participants provided written informed consent.

### 2.2 Session Schedule

Participants completed five training sessions over approximately one week (sessions 1–5, alternating days), plus a follow-up session (Session 6) a median of 29 days after the final training session (mean 29.7, SD 3.4, range 20-39 days; recorded dates available for 35 of 40 participants). BioSemi 64-channel EEG was recorded concurrently with biofeedback on sessions 1, 3, 5, and 6. Sessions 2 and 4 used the biofeedback system only. ERSP analyses use all four BioSemi sessions.

### 2.3 EEG Acquisition

A BioSemi ActiveTwo system recorded 64-channel EEG (active Ag/AgCl electrodes, 10-20 layout, 512 Hz) with bilateral ear references and six ExG channels. Concurrent single-channel biofeedback was acquired via a ProComp+ Infiniti (Thought Technology Ltd.) at the training site (C3-A1 or C4-A2), feeding into EEGer software. The ProComp+ is optically isolated from the BioSemi system. EEGer transmitted TTL triggers to the BioSemi status channel at each reward event onset.

### 2.4 Neurofeedback Protocol

Training used a three-band protocol: reward (SMR 12–15 Hz or Beta 15–18 Hz), inhibit theta (4–7 Hz), and inhibit high beta (22–40 Hz). Reward required simultaneous threshold achievement in all three bands for 500 ms continuously. Reward events consisted of a simultaneous 125 ms auditory beep and picture-grid reveal (both triggered at reward onset; the concurrent visual event occurs within the P2 analysis window). Auto-thresholding recalibrated every 30 seconds, targeting ∼70% reward rate. Each 30-minute session yielded approximately 600–700 reward events. Reward rates did not differ across groups (mean = 618 ± 45 events/session; F(3,36) = 0.29, p = 0.83). Trial counts entering the reward-locked ERP average (§2.6) therefore did not differ by group, eliminating power-asymmetry confounds in the P2 contrast.

### 2.5 Active-Placebo Sham

The sham was developed in collaboration with EEGer Inc. and constitutes an active placebo that eliminates contingency while preserving realistic non-specific effects. Fifteen pre-recorded 3-minute clean EEG segments were assembled in random order to produce a 30-minute sham signal, scaled in real time to match the participant’s broadband EEG amplitude (the fifteen sham segments were amplitude-matched to each other and to the live raw signal before blending). Blinks, muscle artifacts, and electrode noise from the participant’s live EEG were merged into the sham display, making it visually indistinguishable from veridical feedback. All reward thresholds and events were derived from the sham signal. Configuration was password-protected and hidden; group assignment was performed by an unaffiliated researcher. This implementation passes three critical tests: (a) the signal looks alive (artifact merging), (b) the reward rate is plausible (auto-thresholding on EEG-like source), and (c) the operator cannot detect condition (hidden configuration). (a) and (b) are infrastructural design properties; (c) is a design-enforced blinding feature rather than a measured blinding-integrity check, since trainer-guess data were not systematically recorded in the original protocol. To our knowledge, this remains the most rigorous neurofeedback sham in the published literature (see Supplementary S5.3 for sham taxonomy).

### 2.6 Preprocessing and ERSP Computation

Raw BDF files were processed in MNE-Python (Gramfort et al., 2013). Data were high-pass filtered at 0.16 Hz, notch filtered at 60 Hz, and re-referenced to the common average. Reward-locked epochs (−500 to +1500 ms) were baseline-corrected (−100 to 0 ms) and artifact-rejected using an automated statistical procedure (see Supplementary S1 for full details), retaining 94–98% of epochs.

ERSP was computed using Morlet wavelets (3–40 Hz) with single-trial normalization (Grandchamp & Delorme, 2011). The ERD magnitude metric (mean dB in the subject’s reward band within the 200–800 ms post-reward window at C3) served as the primary dependent variable. ERP analysis used the same epochs bandpass-filtered 0.5–30 Hz. Resting-state power spectral density was estimated by Welch’s method.

### 2.7 Statistical Analysis

Between-group ERD differences were tested with a linear mixed-effects model (ERD ∼ Group × Session, random intercept by subject) and planned contrasts (independent t-tests with Cohen’s d, Bayes factors, and FDR correction). Cluster-based permutation tests (Maris & Oostenveld, 2007; 1000 permutations) controlled family-wise error across the time-frequency plane. ERP components (P50, N1, P2) were tested with 4 Group × 4 Session mixed ANOVAs. Resting-state change scores (Session 6 minus Session 1) were compared between groups with FDR-corrected t-tests; a growth curve model tested pre-session baseline trajectories. Full statistical specifications are in Supplementary S2.

Individual alpha frequency (IAF) was estimated as the posterior alpha peak for each resting-state recording. In the reanalysis pipeline, alpha power was computed both in a fixed 8–13 Hz band and in each participant’s individualized band (IAF ± 2 Hz); the primary channel analyses in the text report the original pipeline’s 8–12 Hz alpha band. Growth-curve models were re-fit on the IAF-anchored band as a robustness check. Per-participant responder status was defined by the sign of the within-subject alpha slope across sessions (positive slope = responder).

An exploratory 64-channel topography of resting-state alpha change (section 3.4) was computed from the full BioSemi montage. Eyes-closed pre-session recordings from sessions 1 and 6 were preprocessed with the same minimal pipeline (0.16 Hz high-pass, 60 Hz notch, interpolation of known bad channels, common average reference) across all 64 channels. Power spectral density was estimated per channel by Welch’s method (2 s segments, matching the primary resting-state analysis) with 200 *μ*V peak-to-peak segment rejection; alpha power (8–13 Hz) was integrated per channel and S6-minus-S1 change scores computed. A quality-control ceiling excluded recordings whose maximum absolute per-channel alpha change exceeded 100 *μ*V^2^, retaining 25 of 38 recordings (SMR *n* = 9, Beta *n* = 3, Sham *n* = 13). The pooled SMR-versus-Sham spatial difference was tested with a cluster-permutation test (Maris & Oostenveld, 2007; 2000 permutations, two-tailed) using BioSemi 64-channel electrode adjacency.

## 3. Results

### 3.1 Frequency-Specific, Contingency-Dependent ERD

The frequency crossover (Figure 2) provides the most direct demonstration of specificity: C3 SMR and C3 Beta, trained at the same electrode with reward bands differing by only 3 Hz, produced ERD peaking at different frequencies. Sham showed a flat profile. Any non-specific explanation would predict identical spectral responses; the crossover is difficult to reconcile with non-specific accounts. Per-subject peak ERD frequencies confirmed the separation quantitatively: C3 SMR peaked at 13.8 ± 2.8 Hz, C3 Beta at 16.1 ± 1.8 Hz (d = −0.89; Supplementary S3.6).

Grand-average ERSP heatmaps at C3 (Figure 1; C4 in Supplementary Figure S2) confirmed this pattern across all sessions: active groups produced frequency-specific ERD in the reward band (C3 Beta at 15–18 Hz, C3 SMR and C4 SMR at 12–15 Hz) while sham showed no reward-band ERD despite identical auditory stimulation. C3 Beta and C4 SMR reached FDR-corrected significance; C3 SMR trended in the same direction (see contrasts below). All groups, including sham, produced theta ERS (4–7 Hz, 100–500 ms), likely reflecting a mixture of early auditory-cortical processing and mid-frontal feedback/reward theta (Cavanagh & Frank, 2014); the latter is expected even in sham because sham participants still receive the tone as (placebo) feedback.

**Figure 1.**
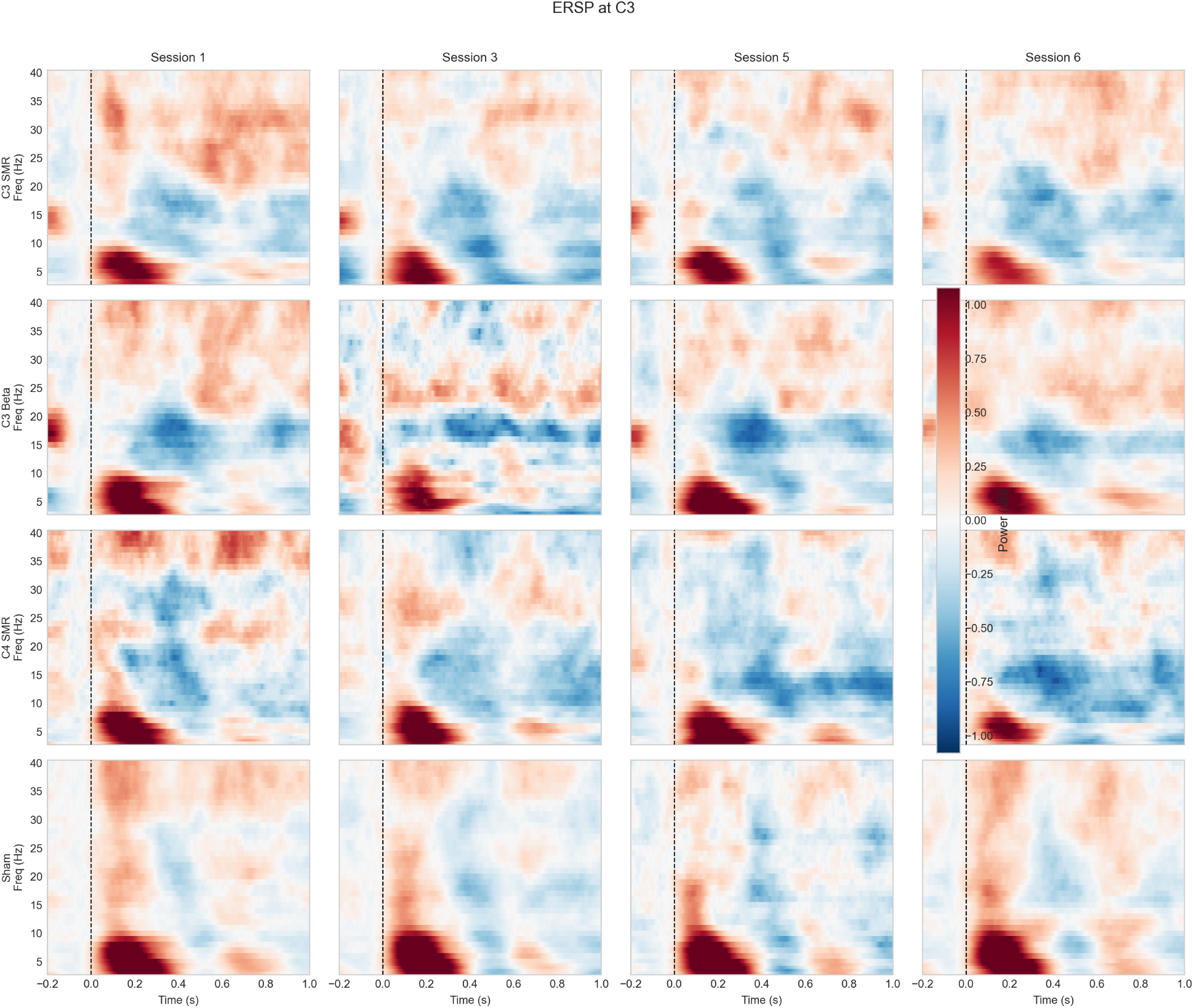
Grand-average ERSP heatmaps at C3 (4 groups × 4 sessions). Active groups show frequency-specific ERD in the reward band; sham shows theta ERS only.

Planned contrasts confirmed a large pooled Active vs Sham effect (d = −1.23, p_adj = 0.001, BF01 = 0.02). The pooled contrast tests the contingency-dependence hypothesis; group-specific contrasts below separately test frequency and site specificity. C3 Beta showed the strongest individual contrast (d = −2.38 vs sham); C4 SMR was significant (d = −1.12); C3 SMR trended (d = −0.80, p_adj = 0.081). An LME model (ERD ∼ Group × Session, random intercept by subject; 157 of 160 possible observations, 3 missing due to excluded sessions) revealed a large Group effect (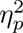 = 0.38) but no Session or Interaction effects, consistent with an ERD that is immediate and stable, present from Session 1 (one-way F(3,36) = 4.72, p = 0.007, 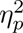 = 0.28; Supplementary §S3.10) and maintained through the retention visit. Paired retention tests (Session 5 vs 6) showed no significant change for any group (Supplementary Table S1).

The scalar ERD contrast between C3 SMR and C4 SMR was small (d = 0.34, 95% CI [−0.74, 1.42], p = 0.49, BF01 = 1.97; Supplementary Table S1); the Bayes factor provides only weak evidence for equivalence at this sample size. However, a cluster permutation test on the Active–Sham time-frequency difference confirmed a significant cluster at C3 (p = 0.002; Supplementary Figure S4) that did not survive correction at C4 (min p = 0.13; Supplementary Figure S5). Site specificity thus appears in the topographic distribution rather than the scalar magnitude. ERD stability across sessions is shown in Supplementary Figure S6. The effect was consistent across individuals, with minimal overlap between active and sham distributions (Supplementary Figure S7). Detailed contrasts, LME coefficients, and retention statistics are in Supplementary S3. Interpretive limits of reward-locked active-vs-sham contrasts arising from the reward criterion’s pre-reward requirement are examined in §4.5; contrasts between active arms are matched on that requirement by design.

### 3.2 ERD–P2 Double Dissociation

The P2 ERP component at C3 showed a large group effect (F(3,33) = 5.02, p = 0.006, 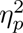 = 0.31; Figure 3), stable across sessions (Group × Session interaction n.s.), paralleling the ERD’s temporal profile. The non-significant interaction is consistent with a between-group site/frequency signature of the operant mechanism rather than a training-induced morphological change, though at these sample sizes we cannot definitively distinguish stable-from-Session-1 from slowly-converging-across-sessions (Session 1 one-way F(3,36) = 0.94, p = 0.43; Supplementary §S3.9). But the groups driving the two effects were different.

**Figure 2.**
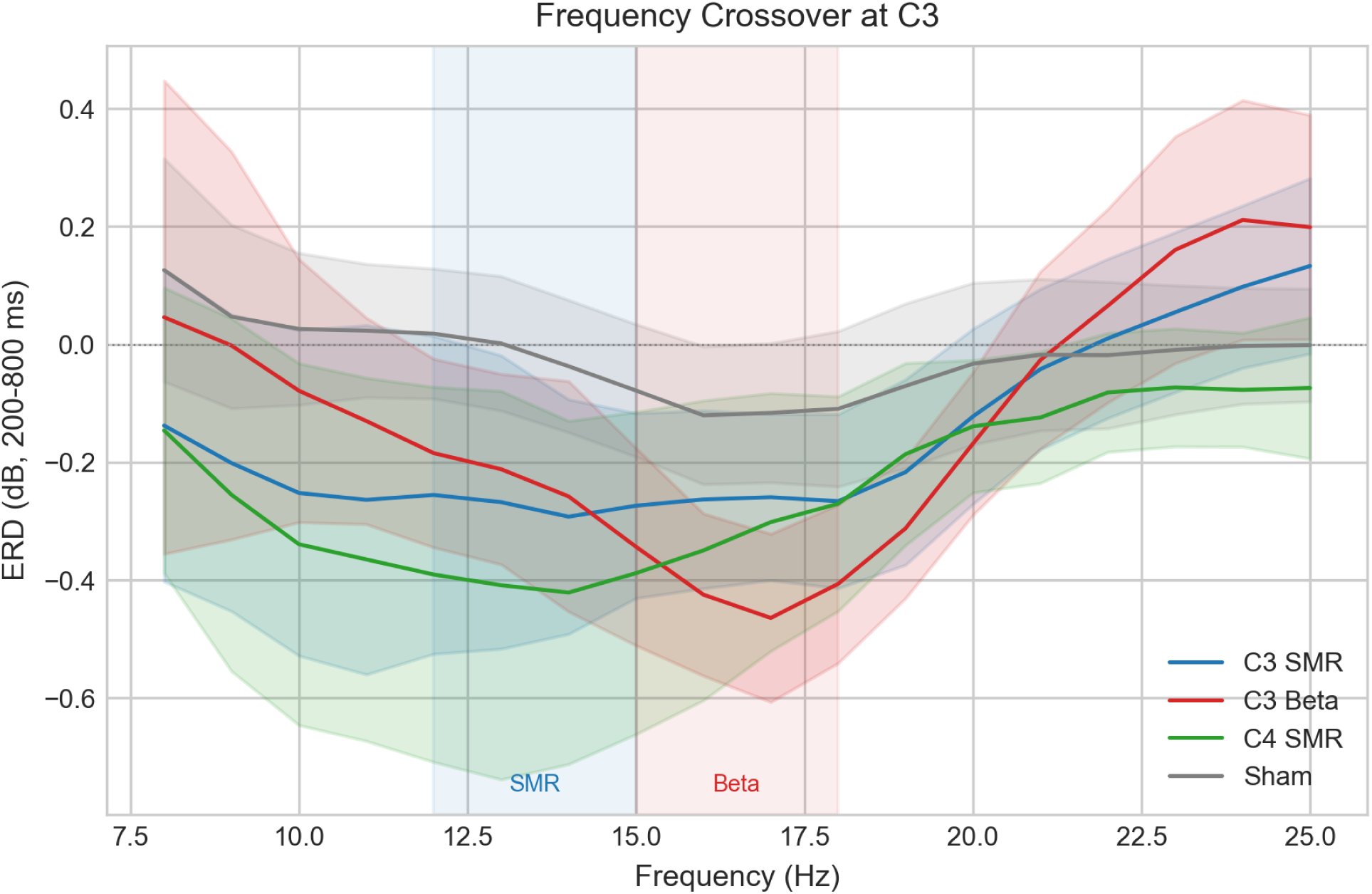
Frequency crossover at C3. C3 SMR ERD peaks at 12–15 Hz; C3 Beta peaks at 15–18 Hz; sham is flat.

**Figure 3.**
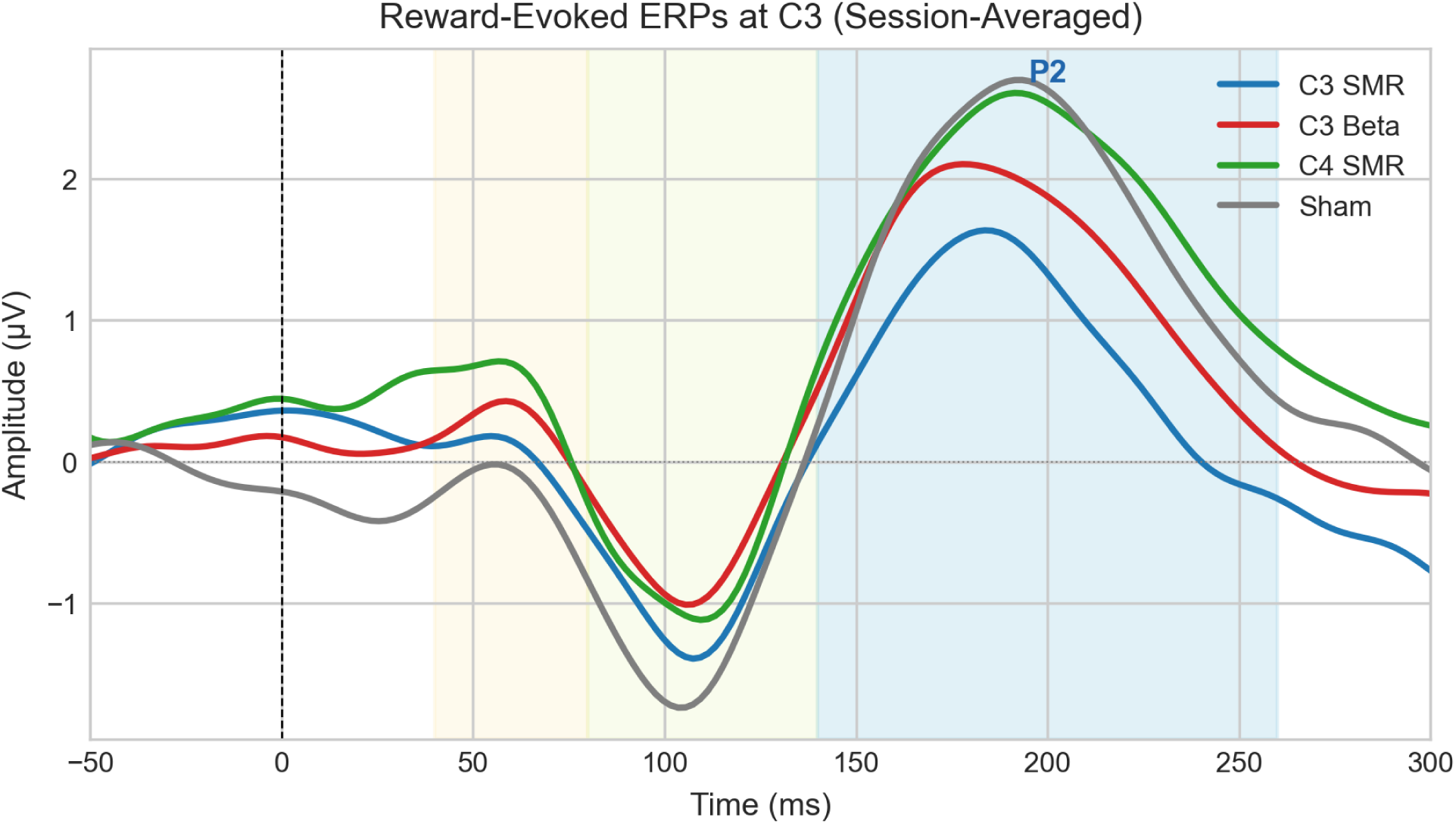
Session-averaged reward-evoked ERPs at C3. The P2 (shaded) is suppressed in C3 SMR relative to the other groups, including sham, the opposite of the ERD pattern (which is strongest in C3 Beta). Full waveforms at C3, C4, Pz in Supplementary Figure S9.

The ERD was strongest in C3 Beta (d = −2.38 vs sham). The P2 was most affected by C3 SMR, which produced a substantially *smaller* mean P2 amplitude at the trained electrode than sham (d = −1.33, BF01 = 0.10, strong Bayesian evidence for a real difference), while C3 Beta (d = −0.27) and C4 SMR (d = 0.19) did not differ from sham. Site-specific suppression was striking: C3 SMR vs C4 SMR d = −1.44 (p_adj = 0.026). The frequency-specificity contrast trended in the *opposite* direction from the ERD (C3 SMR vs C3 Beta d = −1.07, p_adj = 0.079 for the P2; cf. the ERD frequency-specificity contrast p_adj = 0.039, opposite direction). Raw group means at C3: C3 SMR +0.30 µV, C3 Beta +1.43 µV, C4 SMR +1.79 µV, Sham +1.64 µV (n = 37 with complete session data; df = 3, 33). Three participants had missing session-level ERP data (108 S3, 110 S5, 131 S3).

These opposing patterns form a double dissociation: **Beta produces the stronger ERD with a preserved P2; C3 SMR produces weaker ERD with a suppressed P2 at the trained site.** We use the term descriptively; the group-by-measure interaction was not tested directly at these sample sizes. Both are contingency-dependent (absent in sham), but they fractionate spectral power modulation (ERD) from the phasic cortical response to the reward stimulus (P2). Per-subject ERD and P2 amplitudes were uncorrelated (r = −0.06, p = 0.79, n = 24 active; Supplementary S3.7), consistent with distinct processes rather than different views of a single one. Two active subjects had missing Session 6 resting-state data, reducing n to 22 for the resting-state prediction analysis reported in §3.3. P50 was null across all sites; N1 showed a session main effect with no group effect, likely reflecting habituation to the reward tone over repeated sessions. Topographic maps confirming lateralized ERD are in Supplementary Figure S3.

### 3.3 Consolidation: Three-Timescale Plasticity

Three timescales of plasticity were dissociable, each with a different specificity profile.

#### Immediate (trial-locked)

The ERD was stable across sessions (LME interaction n.s.) and retained at the one-month follow-up. Per-subject ERD magnitude during training showed no detectable association with eyes-closed resting-state alpha change at follow-up (trained-site ERD versus trained-channel alpha change: r = 0.28, p = 0.22, n = 22 active subjects), consistent with the dissociation between acute control and durable consolidation developed in §4.4. An earlier version of this analysis reported r = 0.54; that value came from a mislabeled computation (C3 ERD for all groups correlated against 12-15 Hz band change averaged across C3 and C4) and is corrected here.

#### Within-session (minutes)

A single training session transiently increased EC alpha power in all groups (d = 0.5–1.1), including sham at borderline (d = 0.53, p = 0.053). ERD magnitude did not predict within-session shift size (r = −0.09, p = 0.67; Supplementary S4.2), dissociating the operant signal from the transient rebound (Supplementary S4.1).

#### Across-session (days to weeks)

Pre-session EC alpha trajectories diverged by group (Supplementary Figure S10). An LME growth curve on the pipeline’s 8–12 Hz alpha band confirmed SMR-specific accumulation: C3 SMR × Session *β* = 1.44 (p = 0.004), C4 SMR × Session *β* = 1.24 (p = 0.012); C3 Beta and sham showed flat or declining slopes (raw values in Supplementary S4.3, LME coefficients in S4.4). Eyes-closed alpha at follow-up was significantly higher in both SMR groups versus sham (C3 SMR d = 0.97, p_adj = 0.012; C4 SMR d = 0.78, p_adj = 0.028). C3 Beta showed no change (d = −0.002). Eyes-open was null for all groups (Supplementary S4.5–S4.6).

#### IAF-anchored robustness

Mean IAF was similar across groups (descriptive means over all eyes-closed pre-session recordings: C3 SMR 10.1 Hz, C4 SMR 10.2 Hz, C3 Beta 10.1 Hz, Sham 9.9 Hz; no baseline-only test was conducted). Re-running the growth curve on each participant’s individualized alpha band (IAF ± 2 Hz) did not weaken the consolidation result (C3 SMR × Session *β* = 1.21, *p* = 0.006; C4 SMR *β* = 1.10, *p* = 0.013; C3 Beta *β* = 0.04, *p* = 0.92). IAF-anchored S6–S1 effect sizes were large in both SMR arms (C3 SMR *d* = 1.48, C4 SMR *d* = 1.10, C3 Beta *d* = 0.06 vs sham). The consolidation is therefore not an artifact of a fixed band mixing signal and noise across individuals with different peak frequencies.

#### Responder analysis

Classifying each participant by the sign of their within-subject alpha slope (a descriptive split, not a validated responder criterion), 13 of 16 SMR participants (81%) were positive responders (C3 SMR 7/8; C4 SMR 6/8), versus 7 of 16 sham (44%, approximately chance) and 4 of 8 Beta. The direction of change was broadly distributed across individuals rather than concentrated in a few outliers.

#### Individual trajectories

Group-mean EC alpha at C3 (sessions 1/3/5/6, µV^2^) revealed two distinct accumulation patterns in the SMR arms: C4 SMR showed steady climbing (14.1, 15.5, 18.1, 17.3), while C3 SMR was flat during training then showed a pronounced increase at the retention visit (10.0, 10.6, 10.8, 15.1). C3 Beta declined (21.1, 18.3, 18.1, 18.7) and sham was flat (11.9, 10.5, 11.8, 10.5). We report this honestly: the C3 SMR effect is substantially follow-up-weighted, which could reflect either delayed offline consolidation or a single-visit fluctuation; the C4 SMR arm’s smooth climb is harder to attribute to a single visit (see §4.5; Figure 5).

#### IAF as outcome (exploratory)

IAF itself trended upward across sessions in the C3 SMR group only (IAF × Session *β* = +0.15 Hz/session, *p* = 0.088; other groups n.s.), suggesting SMR training may shift the alpha peak frequency; this requires prospective testing.

### 3.4 Exploratory Topography of Consolidation (preliminary)

A 64-channel recompute of resting-state alpha change (S6 minus S1) concentrated the SMR-specific increase over parietal-occipital scalp sites rather than over the trained C3/C4 sensorimotor sites. After QC exclusion of artifact-corrupted recordings (retaining SMR n = 9, Beta n = 3, Sham n = 13), the largest SMR-minus-Sham alpha increases were at posterior sites (P4 Δ = +20.4, P6 +17.4, PO3 +16.9, POz +16.2, Pz +11.7 µV^2^) compared with smaller central effects (C3 Δ = +5.1, C4 +6.0 µV^2^). A spatial cluster-permutation test returned a single posterior/right-parietal cluster (Pz, C2, C4, CP2/CP4, P2/P4) at *p* = 0.082, near-significant but not surviving correction (Supplementary Figure S12).

This result is **exploratory and hypothesis-generating only**: the 64-channel recompute was artifact-prone (heavy QC exclusions reduced the Beta panel to *n* = 3), and the cluster does not survive correction. The confirmatory consolidation claim rests on the C3/C4 channel-level effect reported in §3.3. What the topography adds is a directional hypothesis: the consolidation may be expressed over a posterior alpha network rather than focally at the trained electrode.

### 3.5 Summary of Mechanistic Profiles

The full battery reveals distinct mechanistic profiles (Table 1).

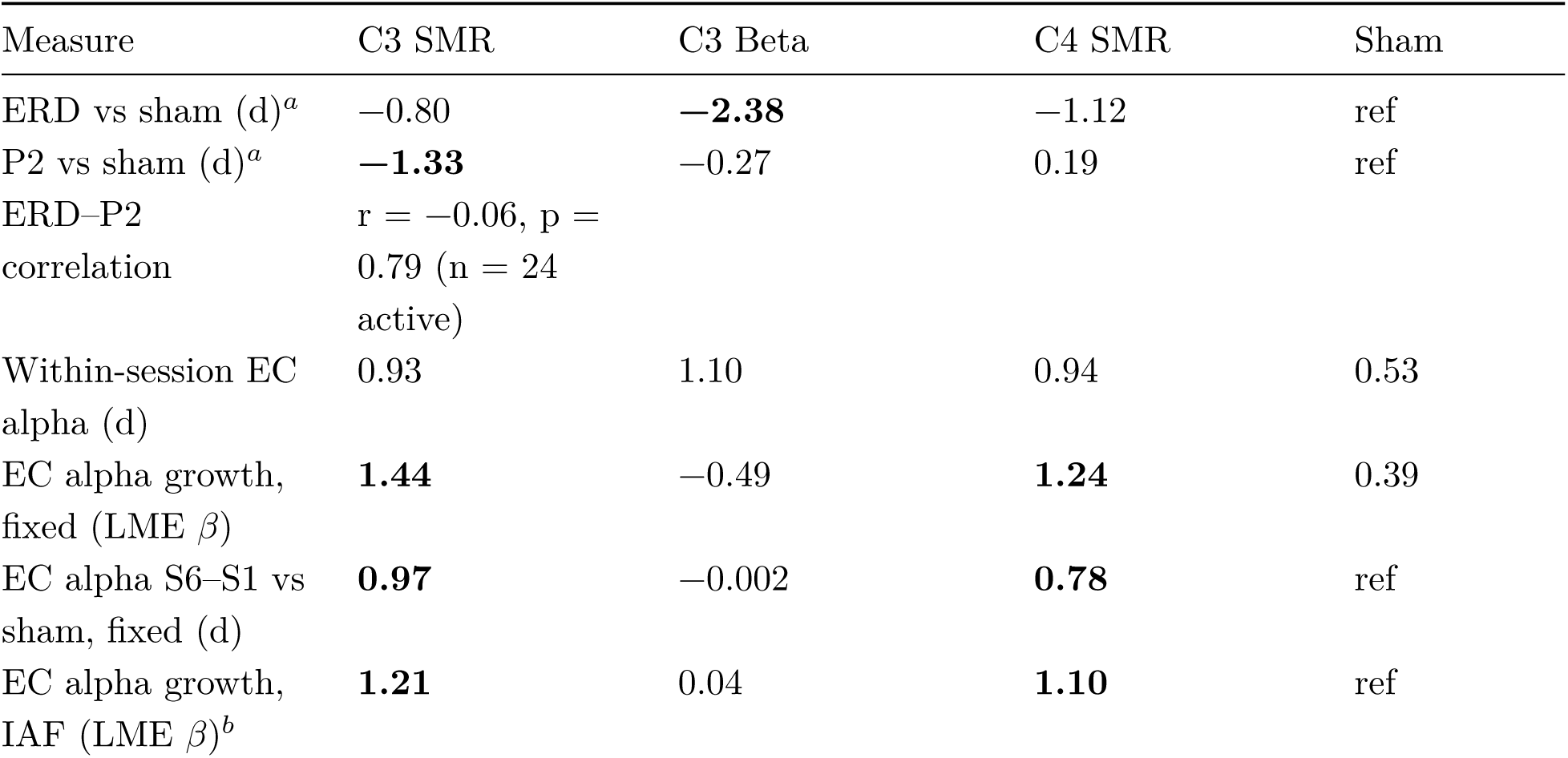
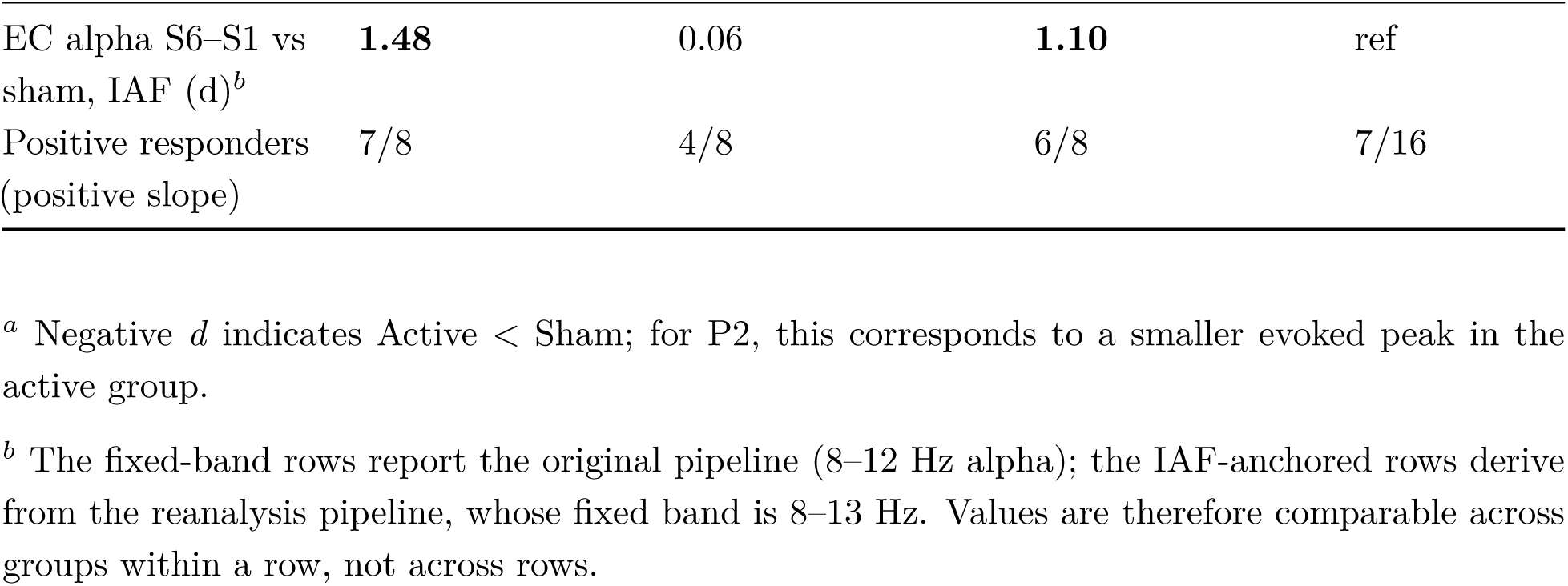

| Measure | C3 SMR | C3 Beta | C4 SMR | Sham |
| --- | --- | --- | --- | --- |
| ERD vs sham (d)<sup>a</sup> | −0.80 | −**2.38** | −1.12 | ref |
| P2 vs sham (d)<sup>a</sup> | −**1.33** | −0.27 | 0.19 | ref |
| ERD–P2 correlation | $r = -0.06$, $p = 0.79$ ($n = 24$ active) | | | |
| Within-session EC alpha (d) | 0.93 | 1.10 | 0.94 | 0.53 |
| EC alpha growth, fixed (LME $\beta$) | **1.44** | −0.49 | **1.24** | 0.39 |
| EC alpha S6–S1 vs sham, fixed (d) | **0.97** | −0.002 | **0.78** | ref |
| EC alpha growth, IAF (LME $\beta$)<sup>b</sup> | **1.21** | 0.04 | **1.10** | ref |
| EC alpha S6–S1 vs sham, IAF (d)<sup>b</sup> | **1.48** | 0.06 | **1.10** | ref |
| Positive responders (positive slope) | 7/8 | 4/8 | 6/8 | 7/16 |

<sup>a</sup> Negative $d$ indicates Active < Sham; for P2, this corresponds to a smaller evoked peak in the active group.
<sup>b</sup> The fixed-band rows report the original pipeline (8–12 Hz alpha); the IAF-anchored rows derive from the reanalysis pipeline, whose fixed band is 8–13 Hz. Values are therefore comparable across groups within a row, not across rows.

The within-session row is the key dissociation: all groups show comparable transient alpha increases (d = 0.53–1.10), yet only the SMR groups consolidate those shifts into cumulative growth. The Beta group shows the *largest* immediate reward-locked ERD yet the *weakest* consolidation, dissociating acute control from durable plasticity. No single measure captures the full mechanistic picture; the implications for the consolidation account are developed in Discussion §4.4.

### 3.6 Sensitivity Analysis

To assess whether results depended on the choice of preprocessing pipeline, we repeated the primary ERD analyses using an ICA-based pipeline (extended Infomax ICA with ICLabel auto-rejection of non-brain components; see Supplementary S1). Trial retention was comparable across pipelines (minimal median = 583, ICA median = 572 per session). Despite similar trial counts, the minimal pipeline produced a somewhat larger Active vs Sham effect (d = −1.23, p = 0.0004) than ICA (d = −1.03, p = 0.0025), and the LME Group effect was larger (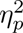 = 0.38 vs 0.26; Supplementary Figure S11). Delorme (2023) benchmarked phase-locked ERP detectability and showed that trial rejection generally reduces detection power rather than improving it; we extend this argument to induced ERD. The minimal pipeline’s statistical artifact rejection preserves more of the reward-locked ERD signal, whereas ICA component removal reduces detection power. All primary findings (frequency specificity, contingency dependence, and the ERD–P2 dissociation) converge across both pipelines.

## 4. Discussion

No single finding in this study would be conclusive at these sample sizes. The evidentiary weight derives from convergence: four independent patterns (SMR-specific resting-state consolidation, frequency-specific ERD, contingency dependence, and the ERD–P2 double dissociation) align within the same 40 subjects under double-blind conditions. Each eliminates a different class of alternative explanation; together, they constrain interpretation to a degree that no single contrast could achieve alone. The most clinically resonant result is the consolidation dissociation: the protocol with the strongest acute control (Beta) fails to consolidate, while the protocol with weaker acute control (SMR) produces durable resting-state growth that persists weeks after training ends.

### 4.1 The Sensory-Operant Dissociation and the Sham

The brain’s reward-locked response contains two distinct signals: a theta-band ERS reflecting a mixture of auditory-cortical processing and mid-frontal feedback/reward theta (Cavanagh & Frank, 2014), present in all groups including sham, and a frequency-specific ERD in the operantly conditioned band that appears only when reward delivery is genuinely contingent on oscillatory power.

The active-placebo sham is critical to this dissociation. The sham produced non-specific effects (theta ERS, behavioral practice) but failed to produce reward-band ERD, with Bayesian evidence favoring the null (BF01 > 3 for all sham reward-band contrasts). This clarifies a current debate about whether sham NF is “partially active” (Supplementary S5.3): it is (the sham produces real sensory-evoked responses), but the contingency-dependent operant mechanism is absent.

Why should reviewers trust this dissociation? The sham signal is pre-recorded EEG with artifact merging from the participant’s live signal, scaled to match real-time amplitude. The reward criterion operates on this sham signal, not on the participant’s actual brain activity. There is no pathway by which the participant’s oscillatory state can influence reward delivery, so reward-band ERD *cannot* emerge through contingency. Theta ERS, by contrast, is a sensory response to the auditory beep itself; it requires only that the participant hear the tone, which all groups do. The theta-present, ERD-absent pattern in sham is exactly what the design predicts. This is the specific signature the Kerson, Sherlin, and Davelaar (2025) framework predicts for non-contingent reinforcement schedules (sensory processing preserved, operant signature absent), and empirically constrains the debate about what “active placebo” sham actually contains.

### 4.2 Frequency Specificity and the Double Dissociation

The frequency crossover provides the most direct test of specificity. As reported in §3.1, two groups trained at the same electrode with reward bands 3 Hz apart produced ERD peaking at different frequencies, a result no non-specific account readily accommodates. This answers Dessy et al. (2020), whose single-reward-frequency SMR design at Cz could not contrast competing trained frequencies within the same paradigm (Supplementary S5.1).

The ERD–P2 dissociation extends the frequency crossover. Both measures are contingency-dependent (absent in sham) but fractionate by group in opposite directions: C3 Beta shows the strongest ERD without changing the P2, while C3 SMR shows moderate ERD with a markedly *suppressed* P2 at the trained electrode. The ERD indexes spectral power change (phase-locked and induced combined); the P2 indexes the phasic evoked response to the reward tone. That C3 SMR training reduces the P2 specifically at the trained site suggests a mechanism that damps the phasic cortical response to the operantly-trained stimulus, a gating profile distinct from the cortical desynchronization engaged by Beta training.

### 4.3 Circuit-Level Interpretation

The double dissociation, combined with the resting-state data, is consistent with a model in which the reward frequency indexes the depth of the circuit engaged, with circuit depth shaping which measures respond and whether effects consolidate. We present this as an interpretive hypothesis; both available direct tests have returned null (below).

SMR at 12–15 Hz is classically tied to thalamocortical relay circuitry implicated in sleep-spindle generation and in sensorimotor gating (Sterman, 1977, 2000; Steriade, McCormick & Sejnowski, 1993; De Gennaro & Ferrara, 2003). If SMR training engages this relay-based thalamocortical mode, a plausible downstream consequence at the trained site is *attenuation* of the phasic evoked response to the operantly-relevant stimulus. The C3 SMR result is consistent with this prediction: moderate ERD at the trained electrode combined with a markedly reduced P2, at that electrode but not at C4. Repeated engagement of this thalamocortical loop could reshape tonic resting-state dynamics. The gating analogy has a logical limit: the reward beep is the operantly-relevant stimulus, not an irrelevant one, so the relay-state interpretation must either confine the gating to the brief reward-delivery window or be understood as a correlate of consolidated training rather than a sleep-spindle-style gating mechanism. An alternative reading that also fits the data is consolidation-driven attentional automatization: as subjects internalize the SMR-to-reward mapping, the reward beep becomes less phasically salient and the evoked P2 diminishes, a mechanism that would co-predict the SMR-specific across-session alpha growth.

Beta at 15–18 Hz is supported by substantial local-cortical mechanisms, including transient beta events arising from pyramidal-cell dendritic integration (Sherman et al., 2016; Shin et al., 2017), with thalamocortical and basal-ganglia contributions also documented (Bonaiuto et al., 2021). A local-cortical ERD without the relay-gating component is consistent with the C3 Beta profile: strongest ERD, preserved P2. But the absence of relay architecture in this account would explain why the local-cortical loop does not propagate into tonic resting-state reorganization in the present data.

The ERD alone is therefore an incomplete index of neurofeedback mechanism. A design measuring only the ERD would rank Beta highest; measuring only the P2 would rank C3 SMR highest in magnitude of change (in the suppression direction). Both measures are needed to differentiate the operant signatures engaged by each protocol.

We emphasize that the present scalp-level data motivate but do not directly test this circuit-level account. Both available direct tests, conducted in response to pre-submission feedback, have returned null. Inter-trial coherence in the reward band at C3 did not differ across groups (F(3,36)

= 1.48, p = 0.24; pooled Active vs Sham *d* = −0.03, BF_01_ = 5.84), and no time-frequency cluster survived permutation testing (C3, 3–20 Hz × 0–1500 ms, 1000 permutations). The ITC–ERD correlation across active subjects was negligible (*r* = −0.06, *p* = 0.79). The reward-band ERD is therefore amplitude-modulated rather than phase-locked. A source-level analysis of reward-locked evoked band-limited power (eLORETA, ico-5 resolution with 20,484 vertices, 1000 permutations for all tests, full-pool noise covariance) likewise found no significant SMR-vs-Beta source difference at any depth weighting (SMR band: 39 clusters, 0 significant, min *p* = 0.785, largest cluster 27 of 20,484 vertices; Beta band: 32 clusters, 0 significant, min *p* = 0.824). Because this analysis projects trial-averaged evoked responses, it tests evoked band-limited source activity rather than the induced single-trial ERD; a single-trial source analysis remains an open extension. An earlier preliminary positive at coarser resolution (ico-4, 100 permutations) did not replicate. Circuit depth thus remains an unconfirmed hypothesis that the sensor-level pattern is consistent with. The contribution of the present data rests on the sensor-level dissociations themselves: frequency-specific ERD, the ERD–P2 double dissociation, and the SMR-specific consolidation of resting alpha. Source-level investigation with intracranial recording or high-density EEG source imaging remains the most direct test of the depth account.

### 4.4 A Consolidation Account

The data reveal a plasticity arc across three timescales. The ERD is the immediate mechanism, present at full strength from Session 1, stable across sessions, maintained at the retention visit. Each training session also produces a transient EC alpha increase (d = 0.5–1.1) in all groups including sham, representing a non-specific post-training rebound. ERD magnitude does not predict within-session shift size (r = −0.09, p = 0.67). No association with the durable resting-state change was detected either (§3.3). This null result argues against the possibility that stronger ERD produces larger transient shifts that then consolidate. Instead, the ERD and the transient rebound appear to be independent processes, with specificity arising at the consolidation step. The LME growth curve shows that only SMR groups’ baselines ratchet upward across sessions (C3 SMR × Session *β* = 1.44, p = 0.004; C4 SMR *β* = 1.24, p = 0.012), while C3 Beta, despite within-session alpha shifts as large as any active group (d = 1.10), shows a *declining* baseline. Each session perturbs the system; the data suggest that only certain circuits consolidate the perturbation.

The consolidation finding is reinforced by three additional analyses. First, IAF-anchored reanalysis confirms it is not a fixed-band artifact (§3.3). Second, 81% of SMR participants are individual responders (§3.3), ruling out an outlier-driven effect. Third, the two SMR arms provide an internal replication across hemispheres.

The arc: **immediate ERD** (trial-locked, stable) then **transient within-session rebound** (universal, non-specific) then **cumulative baseline drift** (SMR-specific, persisting to the one-month follow-up, d = 0.97 vs sham). We propose the following mapping onto Davelaar’s stages, recognizing that our temporal grain (single-trial ERD, across-session growth, multi-week consolidation) differs from Davelaar’s original division: Stage 1 (implicit basal-ganglia-mediated operant learning; Davelaar, 2018) maps onto the immediate ERD; Stage 2 (circuit-level consolidation of repeated perturbation; Davelaar, 2018) maps onto the across-session accumulation seen only in SMR groups; Stage 3 (interoceptive homeostasis; Kerson, Sherlin & Davelaar, 2025) addresses the transition from externally reinforced to self-sustained regulation that the present data cannot test but that the framework predicts. The clinical hypothesis this raises is that the key variable is not session count but *consolidability*, the capacity of each session’s transient shift to persist offline. The protocol with the strongest acute control (Beta) is precisely the one that fails to consolidate; both SMR arms, with weaker acute ERD, produce durable resting-state growth. This dissociation between control and consolidation is independent of the circuit-depth interpretation (§4.3) and stands on the sensor-level data alone. While this sequence is supported by the present dissociations, additional longitudinal work is needed to confirm the causal structure. If confirmed, the model implies that protocol selection should be guided not by session count alone but by the consolidability of the circuit engaged.

### 4.5 Limitations, Future Directions, and Broader Implications

Group sizes were modest (8–16). The consolidation claim rests on the growth-curve LME model and the two-arm SMR internal replication (C3 and C4 SMR both significant), not on any single effect size; the within-group C3 SMR S6–S1 change alone is marginal (*p* ≈ 0.055), which is why the LME and the vs-sham contrasts carry the finding. Effect sizes should be interpreted as upper bounds likely to attenuate in larger samples; the critical finding is the convergence of four independent patterns within the same participants, not the magnitude of any single contrast. The data were collected in 2010–2011 and originally analyzed descriptively (Hill, 2012); the present reanalysis applies a pre-specified modern framework to existing data. The neurofeedback training protocols used fixed frequency bands (12–15 Hz, 15–18 Hz) rather than IAF-anchored bands; IAF-anchored resting-state analyses now confirm the consolidation finding survives individualization (§3.3), and fixed-band training may represent a conservative test, as IAF-anchored protocols should produce stronger effects. Five training sessions address mechanism, not clinical dose-response.

The C3 SMR consolidation effect is follow-up-weighted: the alpha trajectory was flat during training sessions and showed a pronounced increase at the retention visit, which could reflect delayed offline consolidation or a single-visit fluctuation. The C4 SMR arm’s smooth climb mitigates but does not eliminate this concern. Distinguishing the two patterns requires denser follow-up spacing in a prospective design.

The exploratory 64-channel topography (§3.4) is preliminary: the full-scalp recompute was artifact-prone (heavy QC exclusions reduced the Beta panel to n = 3), and the posterior/right-parietal cluster does not survive correction (*p* = 0.082). It is reported as a directional hypothesis, not a confirmed result.

Both direct tests of the circuit-depth hypothesis returned null: the ITC prediction was rejected (§4.3), and source localization (eLORETA, ico-5, 1000 permutations) found no SMR-vs-Beta source difference. This does not invalidate the sensor-level dissociations, which stand independently, but the circuit-depth account remains unconfirmed. Two extensions follow: single-trial ERSP for responder subtypes, and a prospective replication with pre-registered effect sizes and denser follow-up to characterize consolidation trajectories.

A further methodological consideration concerns the pre-reward baseline. Because reward required 500 ms of above-threshold reward-band power immediately before delivery, the −100 to 0 ms reference window is a conditioned state in active groups, and some post-reward decline is expected from regression to the mean in active groups only. A band- and site-matched re-referencing probe confirmed both halves of this concern. Active participants’ pre-reward baselines were elevated relative to the late epoch compared with sham (Hedges *g* = +0.85 to +1.36 across arms), which directly demonstrates that reward delivery was contingent on the trained band in the active arms and not in sham, a manipulation check rarely available in neurofeedback designs. However, re-referencing the ERD window to the late epoch (1200–1500 ms) eliminated the active-sham differences (all |*g*| ≤ 0.23), so at this epoch length the reward-locked ERD cannot be separated from the criterion-conditioned baseline; the pattern persisted when analysis was restricted to trials whose late epoch was uncontaminated by the following reward’s criterion period (next reward at least 2.1 s away). The late-epoch reference is itself imperfect (it can contain the following reward’s criterion period), so a matched pseudo-event control on the continuous recordings, reproducing the selection rule without reward delivery, is required to isolate the reward-specific response and is planned before journal submission. We note that contrasts between active arms (frequency and site specificity, and the ERD-P2 dissociation) compare groups with matched criterion structure and are therefore not affected by this asymmetry.

In response to pre-submission feedback, we tested for a classical Sterman-style post-reinforcement parietal alpha synchronization (Pz/P3/P4, 8–12 Hz, 800–1500 ms post-reward; Supplementary §S3.8) and did not detect one: pooled Active vs Sham d = +0.16, BF01 = 5.20 (moderate evidence for H0), with all three per-group contrasts against sham likewise null (BF01 = 3.30–4.66) and a whole-plane cluster permutation at Pz (3–20 Hz × 0–1500 ms) yielding no significant post-reward cluster. The operant signature in this paradigm therefore appears to be carried by the reward-band ERD itself rather than by a distinct post-reinforcement synchronization burst; whether PRS is genuinely absent in reward-contingent human NF or whether our paradigm masks it (the simultaneous picture-grid reveal dominates the parietal alpha response in this window; Supplementary §S3.8) and Sterman’s original cat/human operant schedules had no such concurrent visual event, is an open question we cannot resolve here.

The reward consisted of a simultaneous auditory beep and picture-grid reveal, so the P2 group effects reflect combined auditory and visual evoked responses rather than a pure auditory P2. This methodological limit, shared with any paradigm delivering multimodal reward at a single time point, means the P2 group differences could partly reflect differential visual-attentional processing; future designs with temporally separated auditory and visual feedback would isolate the modality-specific contribution.

This work provides mechanistic evidence the field needs to move beyond the current efficacy stalemate. The reward-locked ERSP framework provides a principled basis for protocol design: if the mechanism is contingency-dependent, then contingency fidelity, not session count or gamification, is the design variable to test first. The consolidation dissociation adds a second design principle: protocol selection could be guided by consolidability, the capacity of each session’s perturbation to persist offline, rather than by acute within-session control alone, pending prospective confirmation.

## Supporting information

Frequency Specific ERSP Supplemental

## Acknowledgments

Data were collected at the Department of Psychology, University of California, Los Angeles, under IRB approval. The author thanks the original dissertation committee and the participants who contributed their time.

## Data and Code Availability

Analysis code is available at https://github.com/DocSalamandyr/Hill_2026_ERSP-neurofeedback-analysis. Derived data (ERSP matrices, ERP averages, resting-state PSD, and statistical outputs) are deposited at https://doi.org/10.5281/zenodo.19555777. The analysis plan was documented in the project repository prior to re-accessing the raw BDF files (see §2.1); the analysis code sufficient to reproduce all reported results is available at the repository above. Raw EEG recordings (BioSemi BDF) are available from the corresponding author subject to a data use agreement, as the original informed consent did not include provisions for unrestricted public sharing. The consolidation and topography analysis scripts (alpha_consolidation.py, make_figure.py, topography.py, robust_topo.py), the per-subject alpha table (alpha_table.csv), the eLORETA source-replication script (reward_eloreta.py), the Figure 4 estimand audit (fig4_estimand_check.py), and the uniform-permutation source rerun (reward_eloreta_1000.py) are available in the same repository.

**Figure 4.**
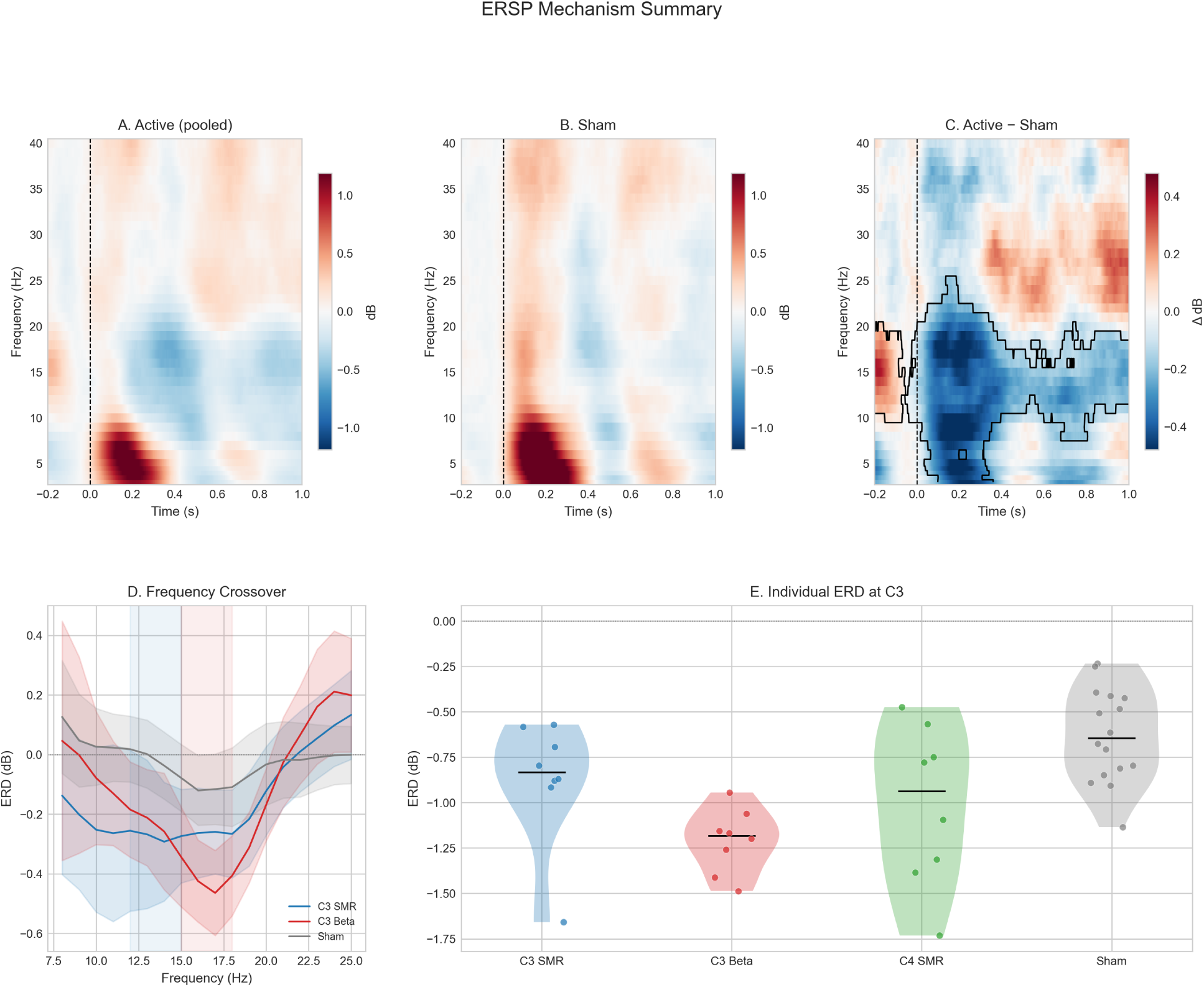
Multi-panel composite summary of the reward-locked ERSP mechanism.

**Figure 5.**
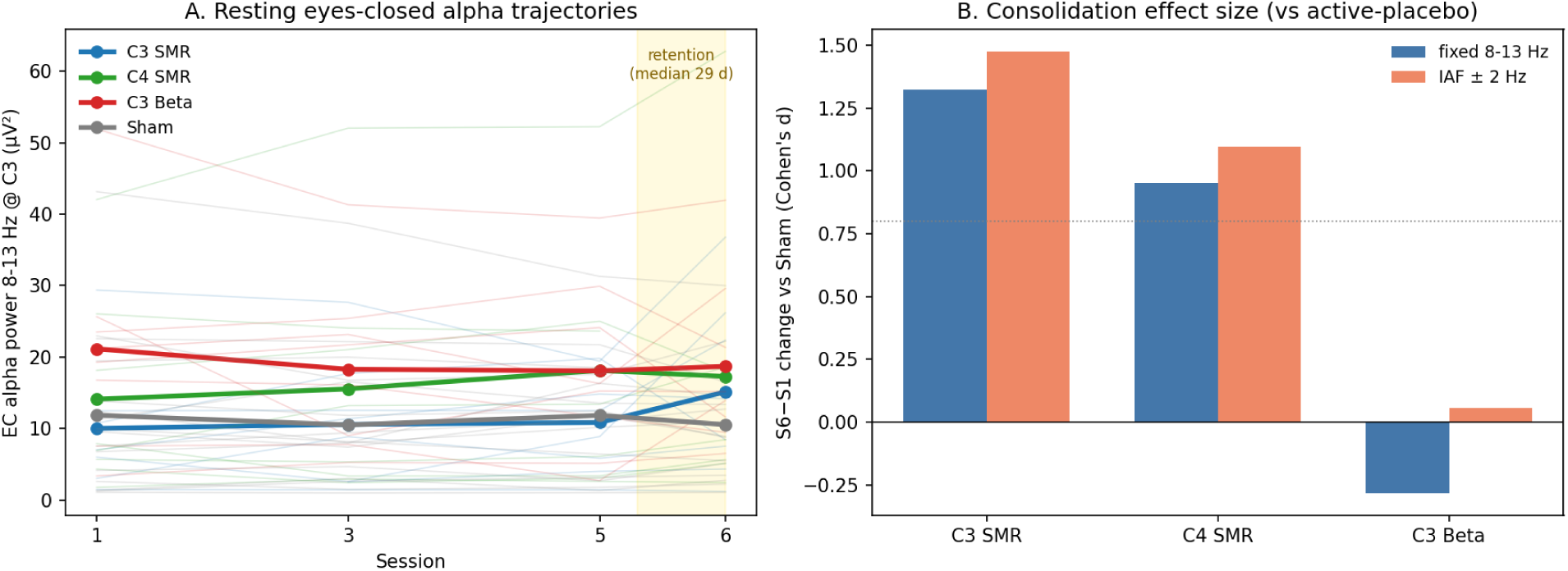
Resting eyes-closed alpha consolidation. (A) Group-mean alpha power (8– 13 Hz) at C3, as computed in the reanalysis pipeline (the channel analyses in the text use the original pipeline’s 8–12 Hz band), across sessions 1, 3, 5, and 6 (retention visit, median 29 days post-training, shaded), with individual-subject trajectories (faint lines). C4 SMR shows steady accumulation across sessions; C3 SMR is flat during training then increases at retention; C3 Beta declines; Sham is flat. (B) S6-minus-S1 consolidation effect sizes (Cohen’s *d* vs active-placebo sham) for fixed 8–13 Hz and IAF ± 2 Hz bands. Both SMR groups show large positive effects in both bands; C3 Beta is near zero. Both panels are computed in the reanalysis pipeline (§2.7); the fixed-band bars are recompute estimates and differ slightly from the published channel-analysis values reported in §3.3 and Table 1.

