## Supplementary material for "Frequency-Specific Operant Learning in Neurofeedback Reveals Distinct Cortical Mechanisms: Evidence from Double-Blind ERSP and ERP Dissociations": Frequency Specific ERSP Supplemental

### Supplementary Materials

Andrew Hill, PhD

Revision, August 2026

#### S1. Preprocessing Pipeline Details

Raw BDF files were imported via MNE-Python 1.9+ (Gramfort et al., 2013, 2014). ExG channels (EXG1–EXG8) were set to EOG/EMG type. Reward events were identified by bit-masking event codes against 0x0100 (256) on the lower 16 bits, capturing both standard codes (256) and variant codes (511, 0x01FF) while stripping BioSemi system flags from higher-order bits. Approximately half of the Session 6 recordings used event code 511 rather than 256, likely reflecting a firmware or configuration difference at the retention visit; the bit-masking approach handles both transparently. All 40 subjects have reward events in Session 6 (range: 368–761 events).

High-pass filtering at 0.16 Hz (zero-phase FIR) was applied, matching the dissertation-era processing. Notch filtering at 60 Hz removed US mains noise. Data were re-referenced to the common average of EEG channels, excluding EOG/EMG channels.

Epochs (−500 to +1500 ms around reward events) were extracted and baseline-corrected (−100 to 0 ms). The shorter baseline window was selected to minimize contamination from adjacent reward events; inter-reward interval analysis confirmed a median IRI of approximately 2.0 s (minimum 0.5 s from the 500 ms reward criterion), meaning longer baselines risk capturing theta ERS from preceding rewards.

**Artifact rejection** used an automated statistical procedure replicating EEGLAB's `pop_autorej` algorithm (Delorme, Sejnowski & Makeig, 2007): (1) epochs exceeding ±1000 µV absolute amplitude were removed; (2) iterative joint-probability rejection identified statistically improbable epochs exceeding 5 standard deviations, with a maximum of 5% of epochs rejected per iteration and adaptive threshold refinement over up to 8 iterations; (3) a final kurtosis check rejected remaining epochs exceeding 6 standard deviations from the mean epoch kurtosis. This approach retains 94–98% of epochs, consistent with Delorme (2023), who demonstrated that trial rejection generally fails to compensate for the associated loss of statistical power. Early/late splits divided each session at the temporal midpoint of the accepted epoch stream.

**Dual-pipeline architecture.** The primary "minimal" mode described above replicates the dissertation-era processing (high-pass filter + statistical artifact rejection only). An alternative "ICA" mode applies the full modern pipeline (0.1 Hz high-pass, RANSAC bad channel detection, extended Infomax ICA with ICLabel auto-rejection of non-brain components at $\geq 0.80$ probability, and 200 µV peak-to-peak epoch rejection). Outputs from both modes are namespaced by preprocessing variant for comparison. Primary results use the minimal pipeline; ICA results are reported as sensitivity analyses.

**ERSP computation.** Morlet wavelets (3–40 Hz; 0.5 Hz steps from 3–5 Hz, 1 Hz steps from 6–40 Hz; 3 cycles at lowest frequency increasing linearly to 12 at highest) with single-trial baseline normalization (Grandchamp & Delorme, 2011). The 3 Hz lower bound captures the full theta band while remaining within the wavelet length constraint imposed by the 2 s epoch. ERP analysis used the same epochs bandpass-filtered 0.5–30 Hz, averaged per subject per session. Resting-state power spectral density was estimated by Welch's method (2 s Hanning windows, 50% overlap, 0.5–45 Hz).

All analysis code is available in the repository (`scripts/pipeline.py`, `tools/`). Python package versions are pinned in `scripts/requirements.txt`.

---

#### S2. Statistical Analysis Details

**Cluster-based permutation tests** (Maris & Oostenveld, 2007) were applied to time-frequency contrasts using 1000 permutations with a cluster-forming threshold of $t = 2.0$, controlling family-wise error across the full time-frequency plane.

**Linear mixed-effects models** tested ERD magnitude (ERD ~ Group × Session) with a random intercept by subject (random slope models did not converge). Fixed effects were Group (4 levels) and Session (4 levels: 1, 3, 5, 6). Partial eta-squared was computed from a parallel mixed ANOVA. Models were implemented via statsmodels MixedLM with pingouin for effect sizes.

**Planned contrasts** used independent-samples t-tests with Cohen's d (95% CI) and Bayes factors (BF01) for evidence quantification. Six planned contrasts (Active pooled vs Sham; C3 SMR vs C3 Beta, frequency specificity; C3 SMR vs C4 SMR, site specificity; and each of the three active groups vs Sham) were corrected using FDR (Benjamini-Hochberg). Bayesian independent-samples t-tests used the Jeffreys scale: $BF01 > 3$ = moderate evidence for absence; $BF01 > 10$ = strong evidence for absence.

**ERP analysis** used 4 Group × 4 Session mixed ANOVA on mean amplitude (µV) for P50 (40–80 ms), N1 (80–140 ms), and P2 (140–260 ms) at C3, C4, and Pz. Post-hoc contrasts on the P2 at C3 (the sole significant component-site combination) used the same contrast structure as the ERD planned contrasts.

**Resting-state analysis** compared Session 6 minus Session 1 change scores (ECPRE, EOPRE) between active groups and sham using independent-samples t-tests with FDR correction across bands and groups. A Group × Condition (EC/EO) mixed ANOVA tested the EC/EO dissociation. Linear regression tested whether per-subject mean ERD during training predicted resting-state power change at follow-up (active subjects only, $n = 22$). Within-session shifts used paired POST minus PRE comparisons averaged across sessions, tested with one-sample t-tests against zero. Pre-session trajectory used an LME growth curve (EC alpha ~ Group × Session, random intercept by subject).

**Retention** was assessed by paired t-tests on ERD between Session 5 and Session 6 per active group with FDR correction.

Software: MNE-Python (Gramfort et al., 2013), statsmodels, pingouin, scipy.

---

#### S3. Extended ERD Results

##### S3.1 Planned Contrast Table (Table S1)

| Contrast | Hedges *g* | 95% CI | *p* | *p*_adj (FDR) | BF01 |
|---|---|---|---|---|---|
| Active (pooled) vs Sham | −1.23 | [−1.94, −0.52] | 0.0004 | 0.0012 | 0.02 |
| C3 Beta vs Sham | −2.38 | [−3.53, −1.24] | <0.001 | 0.0001 | <0.01 |
| C4 SMR vs Sham | −1.12 | [−2.08, −0.16] | 0.0135 | 0.0270 | 0.24 |
| C3 SMR vs Sham | −0.80 | [−1.73, +0.13] | 0.0676 | 0.0812 | 0.73 |
| C3 SMR vs C3 Beta (frequency) | +1.17 | [+0.01, +2.34] | 0.0263 | 0.0394 | 0.37 |
| C3 SMR vs C4 SMR (site) | +0.34 | [−0.74, +1.42] | 0.4883 | 0.4883 | 1.97 |

Note: At n = 8 per group, the default Bayesian prior is diffuse and even moderate true effects ($d \approx 0.5$) can produce BF01 > 3; the site-specificity result (BF01 = 1.97) should therefore be interpreted as weak evidence at best.

Exported deterministically from scripts/export_planned_contrasts.py (session-mean primary ERD at C3; Hedges g with small-sample correction; BH-FDR across the six-test family). Effect sizes reproduce the previously published values exactly; the raw and adjusted p columns and the Bayes factors supersede the earlier table, whose raw column had duplicated the adjusted values.

##### S3.2 LME Omnibus

Linear mixed-effects model (ERD ~ Group × Session, random intercept by subject; 157 observations, 40 subjects): Group main effect $\eta_p^2 = 0.38$; Session main effect $\eta_p^2 = 0.048$ (n.s.); Group × Session interaction $\eta_p^2 = 0.051$ (n.s.). Sham coefficient $\beta = 0.67$ ($p < 0.001$). All Group × Session interaction terms $p > 0.16$.

##### S3.3 Retention Paired Tests

Session 5 vs Session 6 (median 29-day gap): C3 SMR $d = 0.13$ ($p = 0.73$), C3 Beta $d = -0.35$ ($p = 0.35$), C4 SMR $d = -0.10$ ($p = 0.78$). BF01 = 2.01–2.87 (equivocal; compatible with stable maintenance but not evidence of equivalence).

##### S3.4 Cluster Permutation Results

Active (pooled) minus Sham difference maps: C3: 47 clusters tested, 1 significant ($p = 0.002$). C4: 59 clusters tested, 0 significant (minimum $p = 0.13$). The C4 asymmetry reflects both pooling (two-thirds of Active trained at C3) and the thalamocortical model prediction that deeper generators produce more diffuse scalp signatures.

##### S3.5 Within-Session ERD Emergence

Early/late within-session comparisons show the ERD strengthening from the early to the late portion of training sessions (Supplementary Figure S1); the Session-1-only ANOVA (§S3.10) separately establishes that the group effect is present in the first session.

##### S3.6 Peak ERD Frequency

To formally quantify the frequency crossover visible in Figure 2, we extracted per-subject peak ERD frequency at C3 (the frequency showing the strongest desynchronization in the 200–800 ms window, searched within 10–20 Hz). C3 SMR subjects peaked at $13.8 \pm 2.8$ Hz (mean $\pm$ SD); C3 Beta subjects peaked at $16.1 \pm 1.8$ Hz. The 2.3 Hz group difference trended toward significance ($t(14) = -1.89$, $p = 0.080$, $d = -0.89$, BF01 = 0.94). The large effect size and equivocal Bayes factor are consistent with a real separation underpowered at $n = 8$ per group. The direction aligns with the trained frequency bands (12–15 Hz vs 15–18 Hz).

##### S3.7 ERD–P2 Independence

To test whether the ERD and P2 reflect the same underlying process, we correlated per-subject session-averaged ERD magnitude with P2 mean amplitude at C3. Among active subjects ($n = 24$), $r = -0.06$ ($p = 0.79$); across all subjects ($n = 40$), $r = 0.15$ ($p = 0.36$). The absence of correlation supports the characterization of ERD and P2 as dissociable mechanisms, spectral power modulation and phase-locked evoked activity, rather than different measures of a single process.

##### S3.8 Post-Reinforcement Synchronization (null result)

Following pre-submission feedback, we tested for a classical post-reinforcement synchronization (PRS; Wyrwicka & Sterman, 1968; Sterman, 1977, 2000): an alpha-band (8–12 Hz) event-related synchronization in parietal electrodes following reward delivery, in a window strictly after the reward-band ERD (800–1500 ms post-reward). The analysis used the same reward-locked epochs and Grandchamp–Delorme single-trial dB normalization as the primary ERSP pipeline, with baseline $-100$ to 0 ms. Pz served as the primary channel; P3/P4/POz/PO3/PO4 as ROI supporting channels; Fz as a non-parietal control. Pre-specified tests included the pooled Active vs Sham one-tailed contrast, per-group contrasts against sham with FDR across the three active groups, Bayesian evidence via JZS BF01, and a whole-plane cluster permutation at Pz (3–20 Hz × 0–1500 ms, 1000 permutations, cluster-forming $t > 2.0$).

The test was null. Pooled Active vs Sham at Pz: d = +0.16, t(38) = +0.53, p = 0.60, BF01 = 5.20 (moderate evidence favoring H0). Per-group contrasts against sham: C3 SMR d = +0.09 (p_adj = 0.97, BF01 = 4.57), C3 Beta d = +0.38 (p_adj = 0.97, BF01 = 3.30), C4 SMR d = +0.02 (p_adj = 0.97, BF01 = 4.66). The SMR-vs-Beta linear contrast (pre-registered to test the thalamocortical prediction that PRS would load preferentially on SMR-trained groups) likewise yielded moderate evidence for H0 (d = −0.33, BF01 = 3.59). The cluster permutation at Pz produced nine candidate clusters with a minimum p = 0.20, no significant post-reward cluster at any frequency–time location. A frontal control contrast at Fz was similarly null (Active vs Sham BF01 = 5.31).

Descriptively, the Pz alpha scalar was strongly negative in every group (C3 SMR −0.67 dB, C3 Beta −0.55 dB, C4 SMR −0.68 dB, Sham −0.69 dB; each group's one-sample-vs-0 test $p < 0.02$), and the magnitudes at Fz were comparable (≈ −0.7 dB), indicating a diffuse post-reward *alpha desynchronization* rather than the focal parietal synchronization that would constitute PRS. We speculate, consistent with the broader visual-attention alpha-suppression literature (Klimesch, 1999), that the simultaneous picture-grid reveal that accompanies the reward beep in this paradigm dominates the parietal alpha response in this window, a concurrent visual-attention event absent from Sterman's original operant schedules. Because the picture-grid reveal is simultaneous with the reward beep onset, this visual event also falls within the P2 analysis window (140–260 ms); the P2 group effects reported in §3.2 therefore reflect the combined auditory-plus-visual evoked response, a methodological limit shared with any paradigm that delivers multimodal reward feedback at a single time point. The primary ERSP findings at C3/C4 in the 200–800 ms reward-band window are unaffected by this null result. The new pipeline stage used to compute the PRS scalars is documented in `tools/prs.py` and `scripts/pipeline.py --stage prs / --stage prs_stats`; pre-specification and per-branch decision rules are in `audit/PRS_ANALYSIS.md`.

##### S3.9 Session 1 P2 Baseline Test

To assess whether the P2 group effect was present from the first training session, we computed a one-way between-group ANOVA on Session 1 P2 mean amplitude at C3 (N = 40). The test was non-significant: F(3,36) = 0.94, p = 0.43, $\eta_p^2 = 0.07$. Per-group means (µV ± SD): C3 SMR 0.39 ± 0.79 (n = 8), C3 Beta 1.31 ± 0.86 (n = 8), C4 SMR 1.47 ± 0.78 (n = 8), Sham 1.23 ± 2.00 (n = 16). C3 SMR was descriptively the lowest group at Session 1, but the between-group variance was not distinguishable from within-group variance. At these sample sizes the Session 1 test is underpowered to detect anything smaller than a very large effect; the non-significant Group × Session interaction reported in §3.2 remains the primary evidence that the P2 group pattern does not change systematically across sessions.

##### S3.10 Session 1 ERD Baseline Test

To assess whether the ERD group effect was present from the first training session, we computed a one-way between-group ANOVA on Session 1 primary ERD scalar at C3 (N = 40). The test was significant: F(3,36) = 4.72, p = 0.007, $\eta_p^2 = 0.28$. Per-group means (dB ± SD): C3 Beta −1.26 ± 0.32 (n = 8), C3 SMR −0.82 ± 0.55 (n = 8), C4 SMR −0.88 ± 0.38 (n = 8), Sham −0.59 ± 0.40 (n = 16). The group ordering matches the full-session-averaged contrasts reported in §3.1, confirming that the ERD group effect is present from the first session. Combined with the non-significant Group × Session interaction in the LME, this supports the characterization of the ERD as immediate and stable across the training protocol.

---

#### S4. Extended Resting-State Results

##### S4.1 Within-Session Shifts by Group (EC Alpha at C3)

| Group | d | p | n |
|---|---|---|---|
| C3 Beta | 1.10 | 0.017 | 8 |
| C4 SMR | 0.94 | 0.033 | 8 |
| C3 SMR | 0.93 | 0.033 | 8 |
| Sham | 0.53 | 0.053 | 16 |

Within-session shifts at C4 showed a similar pattern (C4 SMR d = 1.05, p = 0.021). Within-session shifts in SMR and beta bands were smaller and non-significant. EO alpha showed no consistent within-session effects.

##### S4.2 ERD Does Not Predict Within-Session Shift

Per-subject ERD magnitude (measured at C3) did not correlate with within-session EC alpha shift size (active subjects: r = −0.09, p = 0.67, n = 24; all subjects: r = −0.07, p = 0.65, n = 40). The within-session alpha rebound is independent of the ERD mechanism, confirming it as a non-specific post-training recovery process.

##### S4.3 Pre-Session Trajectory Raw Values (EC Alpha at C3)

| Group | S1 | S3 | S5 | S6 | S6-S1 d | S6-S1 p |
|---|---|---|---|---|---|---|
| C3 SMR | 9.41 | 9.84 | 9.94 | 14.05 | 0.90 | 0.055 |
| C4 SMR | 13.04 | 14.42 | 17.06 | 16.27 | 0.43 | – |
| C3 Beta | 20.05 | 17.46 | 16.83 | 17.78 | – | – |
| Sham | 10.43 | 9.41 | 10.61 | 9.39 | – | – |

##### S4.4 LME Growth Curve Coefficients

EC alpha ~ Group × Session (random intercept by subject), reference group = C3 Beta.

**At C3**: C3 Beta slope $\beta = -0.49$ (p = 0.155); C3 SMR × Session $\beta = 1.44$ (p = 0.004); C4 SMR × Session $\beta = 1.24$ (p = 0.012); Sham × Session $\beta = 0.39$ (p = 0.351).

**At C4**: C3 SMR × Session $\beta = 0.67$ (p = 0.21); C4 SMR × Session $\beta = 0.48$ (p = 0.36). Directional but non-significant.

##### S4.5 EC vs EO Dissociation

Mixed ANOVA (Group × Condition [EC/EO]) on alpha-band change scores: interaction $\eta_p^2 = 0.147$ (p = 0.149; n = 37 complete cases). Eyes-open resting-state changes showed no significant active-vs-sham effects in any band (all p > 0.19).

##### S4.6 Follow-Up by Band

Eyes-closed alpha (8–12 Hz): C3 SMR d = 0.97 (p_adj = 0.012), C4 SMR d = 0.78 (p_adj = 0.028), C3 Beta d = −0.002 (null). SMR band (12–15 Hz): C3 SMR d = 0.69 (p_adj = 0.106, trending). All other bands (theta, beta) non-significant.

##### S4.7 IAF-Anchored Robustness

Mean individual alpha frequency (IAF) was similar across groups (descriptive means over all eyes-closed pre-session recordings; no baseline-only test was conducted) (C3 Beta 10.08 Hz, SD 0.72; C3 SMR 10.12 Hz, SD 1.01; C4 SMR 10.23 Hz, SD 0.81; Sham 9.94 Hz, SD 1.02). Growth-curve models were re-fit using each participant's individualized alpha band (IAF ± 2 Hz); recordings without an available IAF estimate were excluded, reducing the IAF-anchored dataset to $n_{\text{obs}} = 143$ (versus 153 for the fixed 8–13 Hz band).

**Table S2. IAF-anchored growth-curve coefficients (EC alpha ~ Group × Session, random intercept by subject).**

| Channel | Group × Session | $\beta$ | $p$ |
|---|---|---|---|
| C3 | C3 SMR × Session | +1.212 | 0.0060 |
| C3 | C4 SMR × Session | +1.096 | 0.0133 |
| C3 | C3 Beta × Session | +0.040 | 0.9249 |
| C4 | C3 SMR × Session | +0.614 | 0.1827 |
| C4 | C4 SMR × Session | +0.459 | 0.3214 |
| C4 | C3 Beta × Session | +0.302 | 0.5008 |

IAF-anchored S6-minus-S1 Cohen's $d$ versus sham: at C3, C3 SMR $d = +1.48$, C4 SMR $d = +1.10$, C3 Beta $d = +0.06$; at C4, C3 SMR $d = +0.69$, C4 SMR $d = +0.71$, C3 Beta $d = +0.38$.

Individualization preserves the consolidation result at C3 and leaves the C4 pattern directionally similar but non-significant, consistent with the fixed-band findings (S4.4).

##### S4.8 Responder Classification and Individual Trajectories

Classifying each participant by the sign of their within-subject OLS alpha slope across sessions (positive slope = responder), 13 of 16 SMR participants (81%) were positive responders (C3 SMR 7/8; C4 SMR 6/8), versus 7 of 16 sham (44%, approximately chance) and 4 of 8 C3 Beta. The group effect is not carried by a few outliers.

Group-mean eyes-closed alpha power at C3 ($\mu V^2$, fixed 8–13 Hz) across sessions 1, 3, 5, and 6:

| Group | S1 | S3 | S5 | S6 |
|---|---|---|---|---|
| C3 SMR | 10.00 | 10.57 | 10.83 | 15.11 |
| C4 SMR | 14.09 | 15.53 | 18.11 | 17.28 |
| C3 Beta | 21.14 | 18.27 | 18.06 | 18.68 |
| Sham | 11.86 | 10.49 | 11.84 | 10.53 |

Two distinct accumulation patterns emerge in the SMR arms: C4 SMR shows steady climbing across sessions, while C3 SMR is flat during training then shows a pronounced increase at the retention visit. C3 Beta declines from a high baseline; Sham is flat. The C3 SMR effect is substantially follow-up-weighted, which could reflect either delayed offline consolidation or a single-visit fluctuation; the C4 SMR arm's smooth climb is harder to attribute to a single visit.

**IAF as outcome (exploratory).** IAF itself was modeled as a dependent variable (IAF ~ Group × Session, random intercept by subject; $n_{\text{obs}} = 143$). C3 SMR showed a non-significant upward trend (C3 SMR × Session $\beta = +0.147$ Hz/session, $p = 0.0882$); C4 SMR ($\beta = -0.021$, $p = 0.8077$) and C3 Beta ($\beta = -0.031$, $p = 0.7131$) showed no trend. This suggests SMR training may shift the alpha peak frequency, but the result requires prospective testing.

##### S4.9 Exploratory 64-Channel Topography of Resting Alpha Change

Eyes-closed pre-session recordings from sessions 1 and 6 were preprocessed across all 64 BioSemi channels with the same minimal pipeline (0.16 Hz high-pass, 60 Hz notch, interpolation of known bad channels, common average reference). Per-channel alpha power (8–13 Hz) was estimated by Welch's method (2 s segments) with 200 $\mu$V peak-to-peak segment rejection, and S6-minus-S1 change scores were computed per participant. A quality-control ceiling excluded recordings whose maximum absolute per-channel alpha change exceeded 100 $\mu V^2$, retaining 25 of 38 recordings (SMR $n = 9$, Beta $n = 3$, Sham $n = 13$; 13 recordings dropped as artifact outliers). The pooled SMR-versus-Sham difference was tested with a spatial cluster-permutation test (2000 permutations, two-tailed) using BioSemi 64-channel electrode adjacency (Maris & Oostenveld, 2007).

**Results.** The cluster-permutation test returned 2 clusters, 0 significant at $p < 0.05$; the smallest cluster $p = 0.0815$, comprising channels Pz, C2, C4, CP4, CP2, P2, and P4.

Top SMR-minus-Sham alpha-change channels ($\mu V^2$, QC-filtered):

| Channel | $\Delta$ ($\mu V^2$) |
|---|---|
| P4 | +20.41 |
| P6 | +17.40 |
| PO3 | +16.87 |
| POz | +16.23 |
| PO4 | +15.50 |
| P2 | +15.20 |
| P3 | +13.26 |
| Pz | +11.67 |
| PO7 | +9.50 |
| P8 | +8.95 |

Central electrode effects were smaller: C3 $\Delta = +5.07$, C4 $\Delta = +6.00$, Pz $\Delta = +11.67$ $\mu V^2$.

This analysis is **exploratory and hypothesis-generating only**. The 64-channel recompute was artifact-prone (heavy QC exclusions reduced the Beta panel to $n = 3$), and the cluster does not survive correction. The confirmatory consolidation claim rests on the C3/C4 channel-level effect reported in the manuscript (section 3.3). The topography adds a directional hypothesis: the consolidation may be expressed over a posterior alpha network rather than focally at the trained electrode (Supplementary Figure S12).

---

#### S5. Extended Literature Context

##### S5.1 The ERD/ERS Framework and Physiological Interpretation

The event-related desynchronization (ERD) and event-related synchronization (ERS) framework, formalized by Pfurtscheller and Lopes da Silva (1999), provides a conceptually grounded approach to understanding transient changes in oscillatory power. Operationally, ERD is a transient decrease in band-limited power; the classical interpretation attributes this decrease to reduced local synchrony of the underlying neural population (Pfurtscheller & Lopes da Silva, 1999).

##### S5.2 Phase-Resetting Versus Additive Models

A fundamental debate concerns whether event-related power changes reflect phase-reset in ongoing oscillations or evoked additive responses (Makeig et al., 2002; Klimesch et al., 2007). Inter-trial coherence (ITC) distinguishes these: high ITC indicates phase-resetting; low ITC suggests additive mechanisms. For operant conditioning, this distinction is meaningful: amplitude-based learning should produce low ITC; phase-alignment learning should produce high ITC.

##### S5.3 The Sham Taxonomy

The challenge of credible sham conditions deserves expanded consideration. Schönenberg et al. (2017) identified several sham classes: pseudorandom/yoked, reversed contingency, orthogonal feedback, and intermittent real feedback. Each has limitations. The active-placebo approach (Ros et al., 2020) provides genuine real-time EEG feedback with critical contingency features hidden. The present study's implementation is detailed in Methods §2.5.

##### S5.4 Theta Oscillations in Learning and Reward Processing

Theta oscillations (4–7 Hz) are associated with memory formation, attentional allocation, and reward processing (Buzsáki & Draguhn, 2004; Cavanagh et al., 2012; Cohen et al., 2007). In the neurofeedback context, theta ERS following the reward beep likely reflects universal sensory-cognitive registration of the auditory stimulus and initiation of reinforcement processes, occurring regardless of contingency.

##### S5.5 The Statistical Modernization Argument

Classical neurofeedback research employed approaches now recognized as statistically fragile. Modern practice uses cluster-based permutation testing to control family-wise error, Bayesian inference to quantify evidence for and against hypotheses, mixed-effects models to capture within-subject variability, and standardized effect sizes for all contrasts. This study adopts a transparency-first reanalysis approach: analytical decisions were documented before BDFs were re-accessed, and the analysis plan is posted to OSF at submission.

---

#### S6. Sensitivity Analysis: Minimal vs ICA Preprocessing

To assess robustness to preprocessing choices, all primary analyses were repeated using an alternative ICA-based pipeline (0.1 Hz high-pass, RANSAC bad-channel detection, extended Infomax ICA with ICLabel auto-rejection of non-brain components at $\geq 0.80$ probability, 200 µV peak-to-peak epoch rejection). Both pipelines processed all 40 subjects.

**Trial retention.** The two pipelines retained comparable trial counts (minimal: median = 583, mean = 516; ICA: median = 572, mean = 521 per session; Figure S11, left panel). Individual sessions showed scattered discrepancies; ICA occasionally rejected substantially more epochs when component decomposition was unstable, but overall yields were within 1% on average.

**ERD effect sizes.** The pooled Active vs Sham contrast was significant under both pipelines: minimal $d = -1.23$ ($p = 0.0004$), ICA $d = -1.03$ ($p = 0.0025$). The LME Group main effect was $\eta_p^2 = 0.38$ (minimal) vs $\eta_p^2 = 0.26$ (ICA). Per-group ERD distributions (Figure S11, right panel) show that the minimal pipeline consistently produces slightly larger effect magnitudes across all active groups.

**Interpretation.** The minimal pipeline's advantage in effect size, despite comparable trial counts, reflects signal preservation rather than trial quantity. ICA component removal can attenuate reward-locked ERD signal mixed into components classified as non-brain (eye, muscle, channel noise). The statistical artifact rejection in the minimal pipeline removes entire epochs with extreme characteristics but preserves the signal in retained epochs. This is consistent with Delorme (2023), who demonstrated that aggressive trial rejection fails to compensate for associated power losses. The convergence of all primary findings across both pipelines confirms that the results are not an artifact of the preprocessing approach.

---

#### S7. Source-Level Analysis (eLORETA, Evoked Band-Limited Power)

To directly test whether SMR and Beta training engage spatially distinct cortical sources, a reward-locked eLORETA source analysis was conducted in response to pre-submission feedback.

**Methods.** An ico-5 source space (20,484 vertices) was constructed. Reward-locked evoked responses from all active subjects (SMR $n = 16$, Beta $n = 8$; 64 channels) were projected to source space using eLORETA inverse solutions. A full-pool noise covariance matrix was estimated from 9,282,840 baseline samples across 24 subjects. Cluster-permutation tests (1000 permutations for all four tests) compared SMR-versus-Beta source activation in both the SMR band (12–15 Hz) and the Beta band (15–18 Hz), at two depth weightings (0.8, standard; 0.0, no depth weighting). An earlier run used 1000 permutations for one test and 500 for the other three; the uniform rerun reported here supersedes it, with the same conclusion.

**Results.** No significant source-level difference was found at any depth weighting or frequency band.

| Band | Depth | Clusters | Significant | Min $p$ | Largest cluster |
|---|---|---|---|---|---|
| SMR (12–15 Hz) | 0.8 | 39 | 0 | 0.7850 | 27 vertices |
| Beta (15–18 Hz) | 0.8 | 32 | 0 | 0.8240 | 18 vertices |
| SMR (12–15 Hz) | 0.0 | 39 | 0 | 0.7850 | 27 vertices |
| Beta (15–18 Hz) | 0.0 | 32 | 0 | 0.8240 | 18 vertices |

Because the inverse solution is applied to trial-averaged evoked responses, this analysis tests evoked band-limited source activity rather than induced single-trial ERD.

The largest cluster at any setting comprised 27 of 20,484 vertices. An earlier preliminary analysis at coarser resolution (ico-4, 100 permutations) had produced a nominally significant cluster; that result does not replicate at the present resolution and permutation count and is superseded. The null result at full resolution, with a properly estimated noise covariance and 1000 permutations, indicates that the reward-locked ERD group differences observed at the sensor level do not map onto a localizable source-space contrast between SMR and Beta training at the resolution afforded by 64-channel EEG.

---

#### Supplementary Figures

- **Figure S1**: Within-session ERD learning curves (early vs late)
- **Figure S2**: ERSP heatmap grid at C4 (4 groups × 4 sessions)
- **Figure S3**: 64-channel topographic ERD maps
- **Figure S4**: Active–Sham difference ERSP at C3 with cluster permutation contours
- **Figure S5**: Active–Sham difference ERSP at C4
- **Figure S6**: ERD magnitude by Group × Session line plot
- **Figure S7**: Individual-subject ERD violin plots
- **Figure S8**: Frequency crossover at C4
- **Figure S9**: Full reward-evoked ERP waveforms at C3, C4, Pz (Sessions 1 and 5)
- **Figure S10**: Pre-session EC alpha growth curve at C3 (all groups across sessions)
- **Figure S11**: Sensitivity analysis: minimal vs ICA preprocessing. Left: trial retention scatter (each point = one subject x session). Right: per-group ERD distributions under both pipelines. The minimal pipeline produces slightly larger effect sizes despite comparable trial counts, consistent with signal preservation over aggressive component removal.
- **Figure S12**: Exploratory 64-channel topography of resting eyes-closed alpha change (S6 minus S1), QC-filtered. Four topographic maps: SMR (n = 9), Beta (n = 3), Sham (n = 13), and the SMR-minus-Sham difference. The SMR group and the difference map show a broad posterior-parietal increase concentrated over right parietal-occipital scalp sites.

#### Supplementary Figures

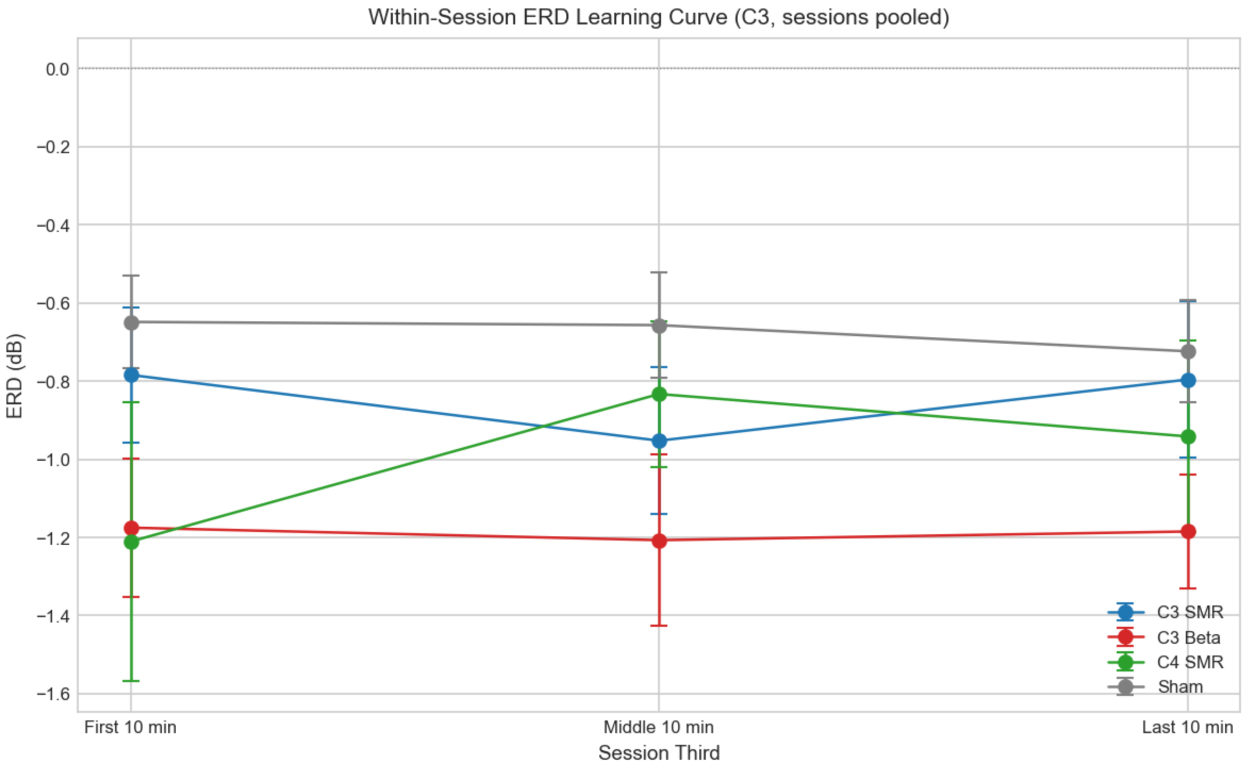


Figure 1: **Figure S1.** Within-session ERD learning curves (early vs late).

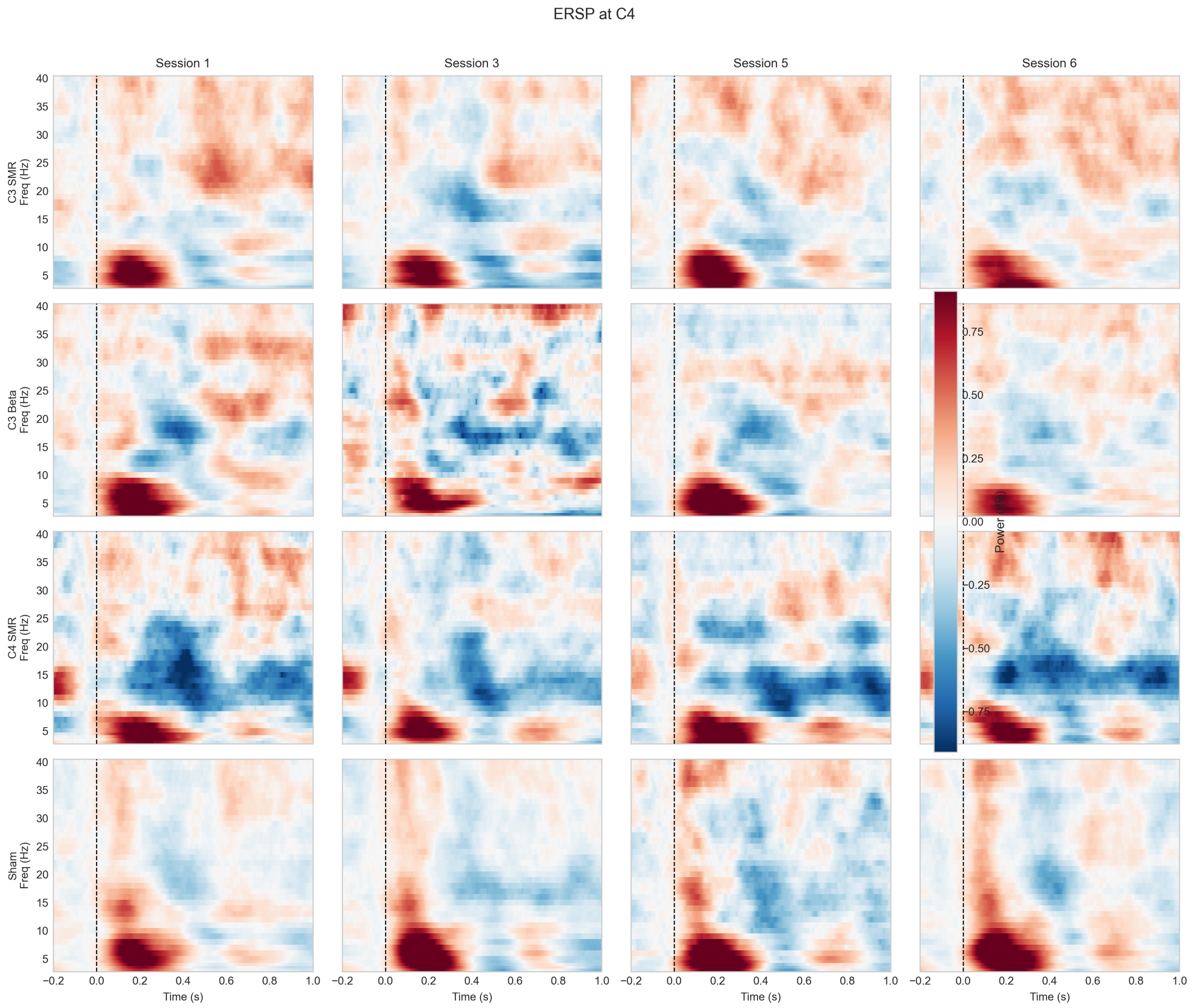


Figure 2: **Figure S2.** Grand-average ERSP heatmaps at C4 (4 groups × 4 sessions).

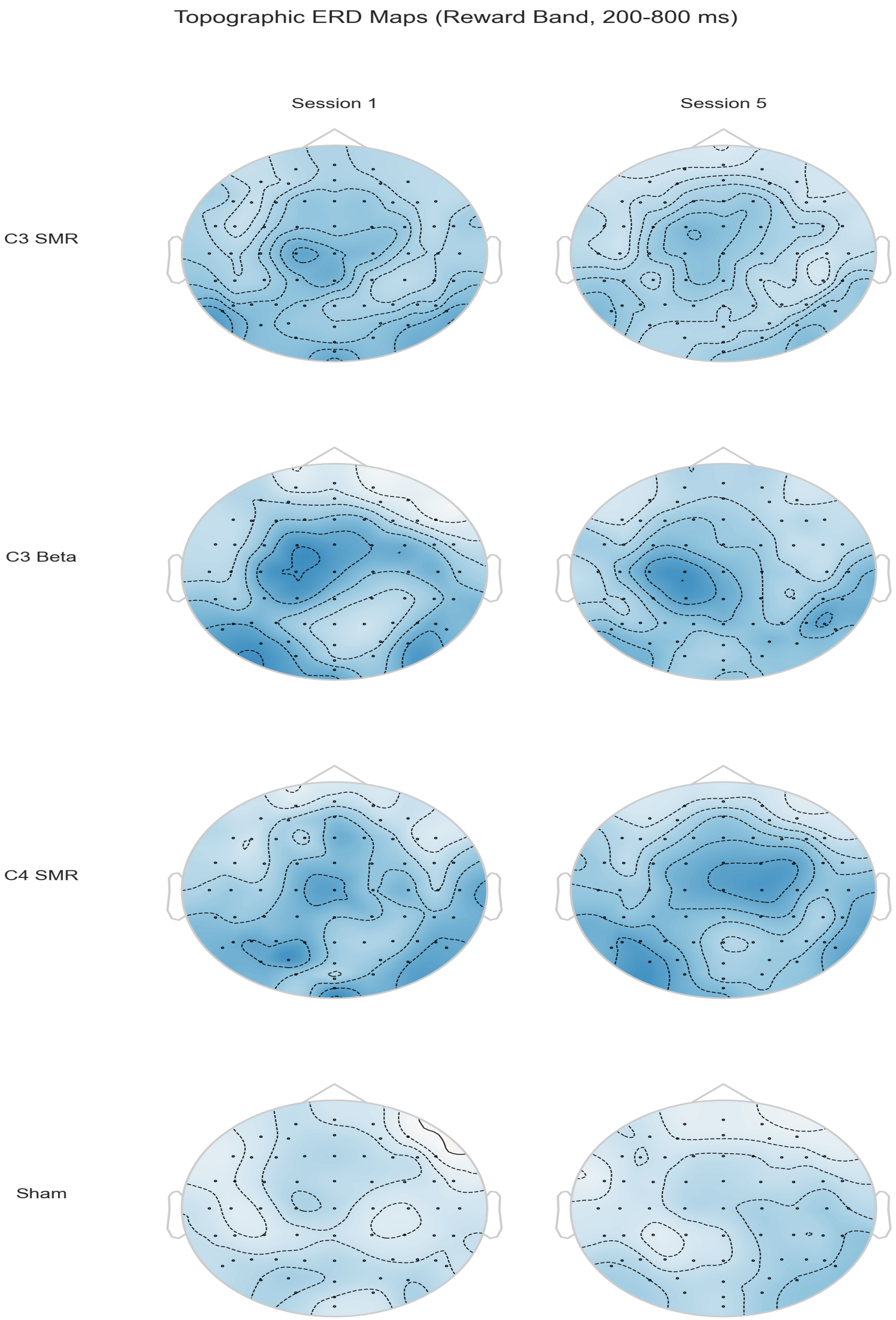


Figure 3: **Figure S3.** 64-channel topographic ERD maps showing lateralized desynchronization.

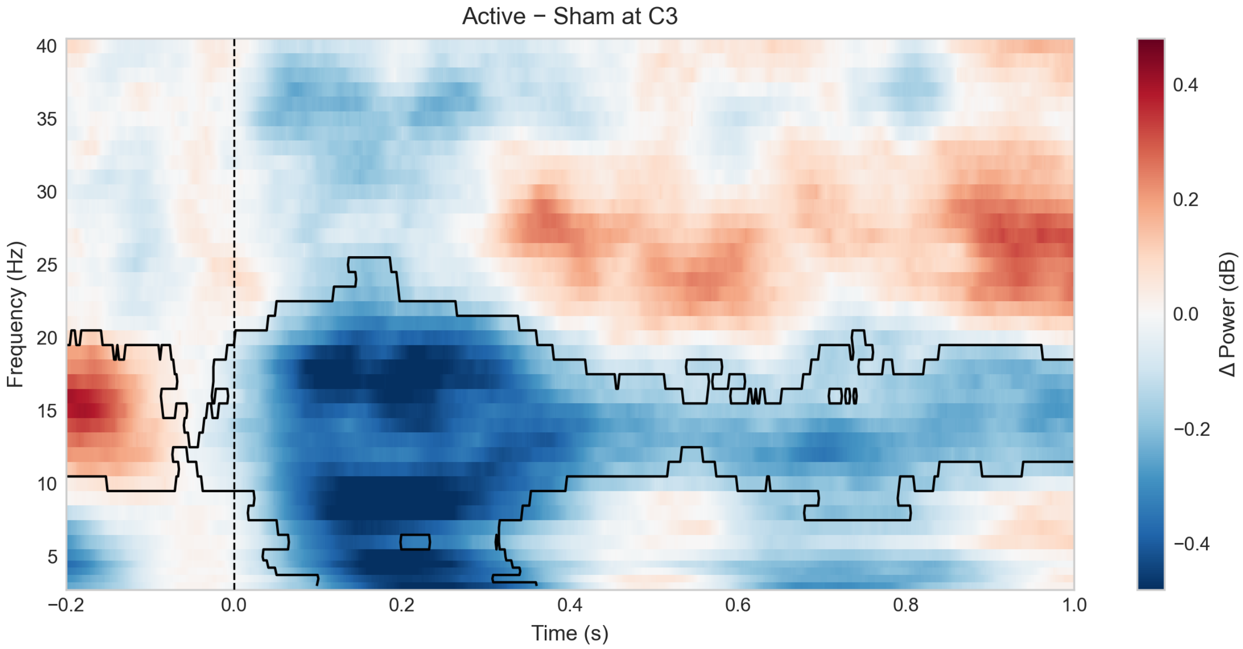


Figure 4: **Figure S4.** Active–Sham difference ERSP at C3 with cluster permutation contours (p = 0.002).

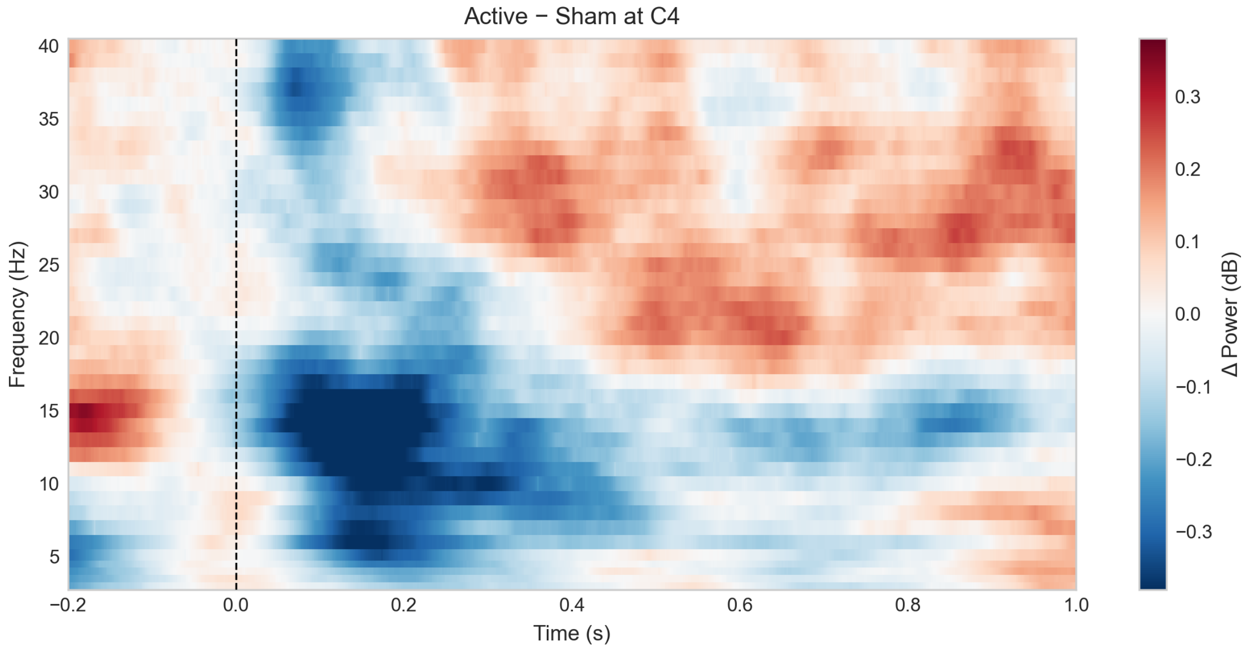


Figure 5: **Figure S5.** Active–Sham difference ERSP at C4 (no clusters survived correction; min p = 0.13).

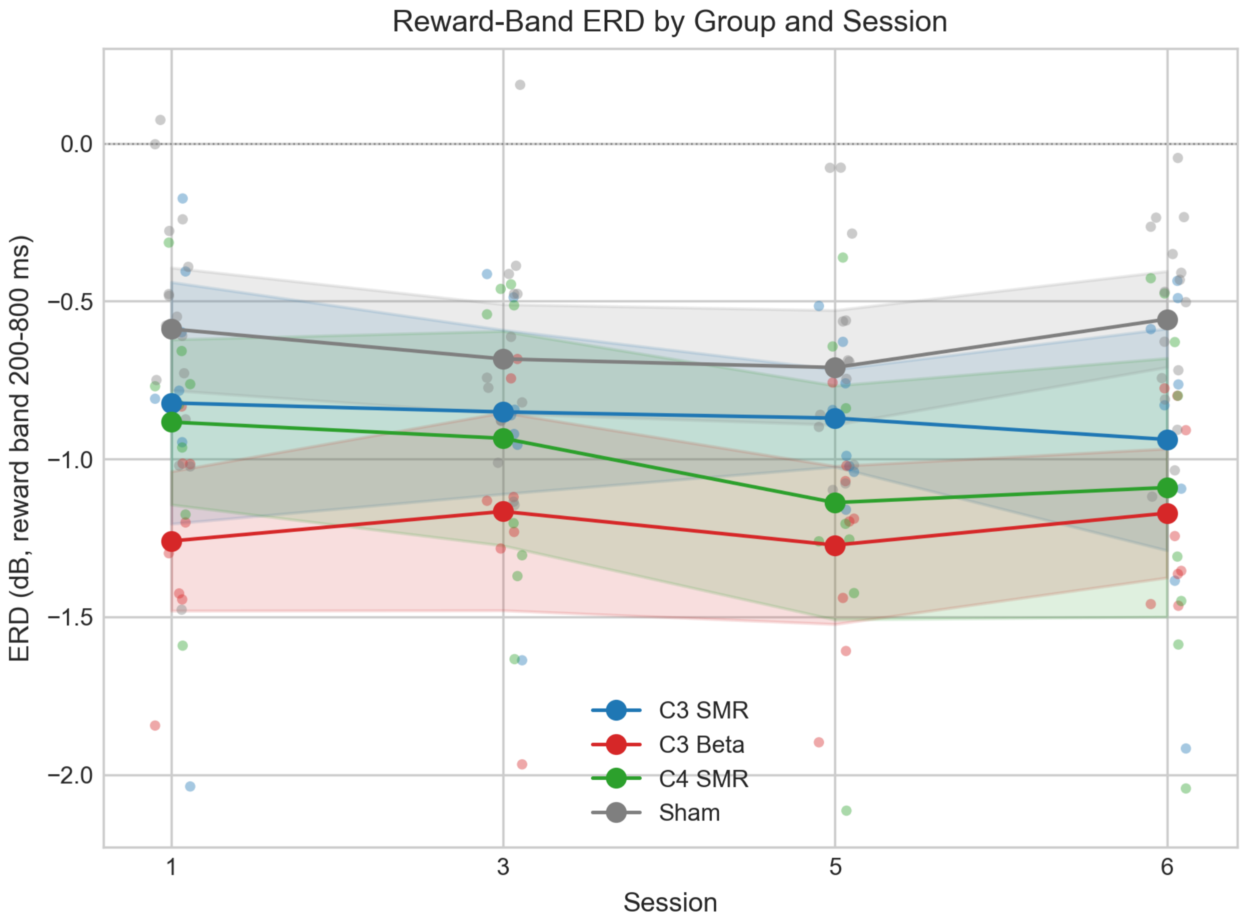


Figure 6: **Figure S6.** ERD magnitude by Group × Session. The ERD is immediate and stable.

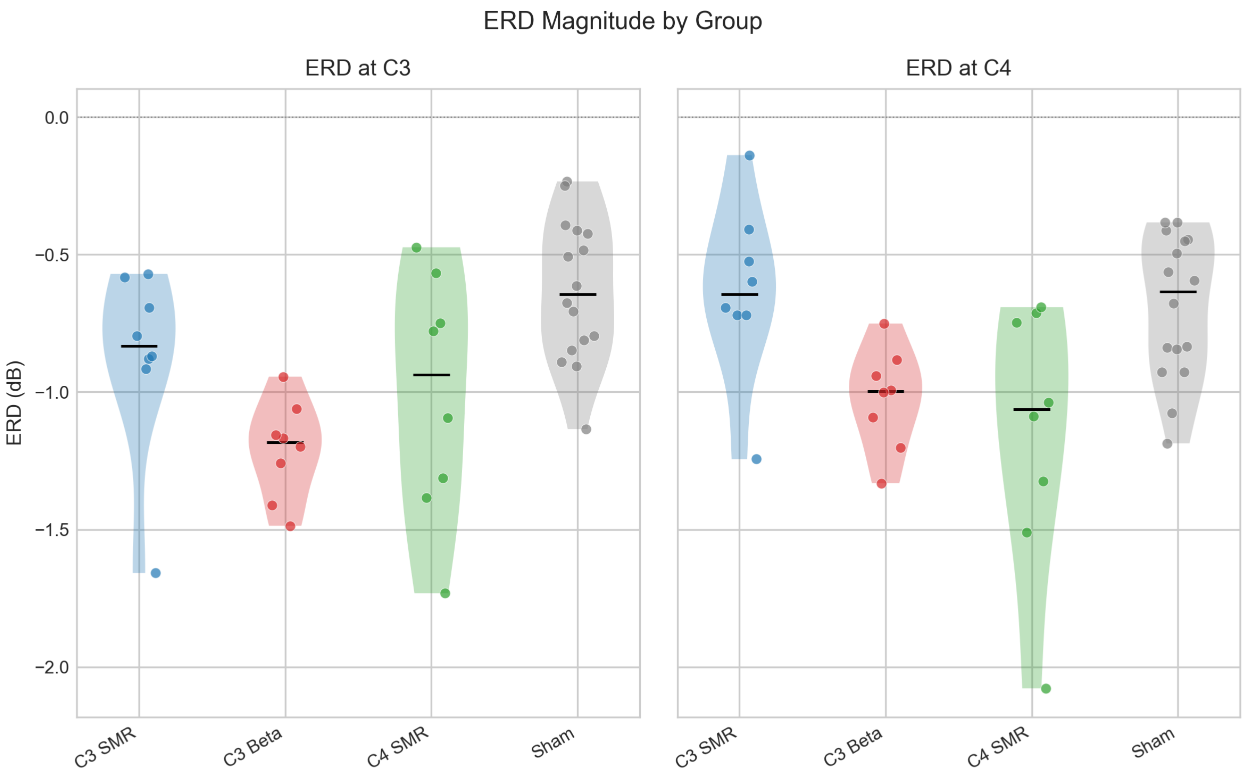


Figure 7: **Figure S7.** Individual-subject ERD violin plots with strip overlay.

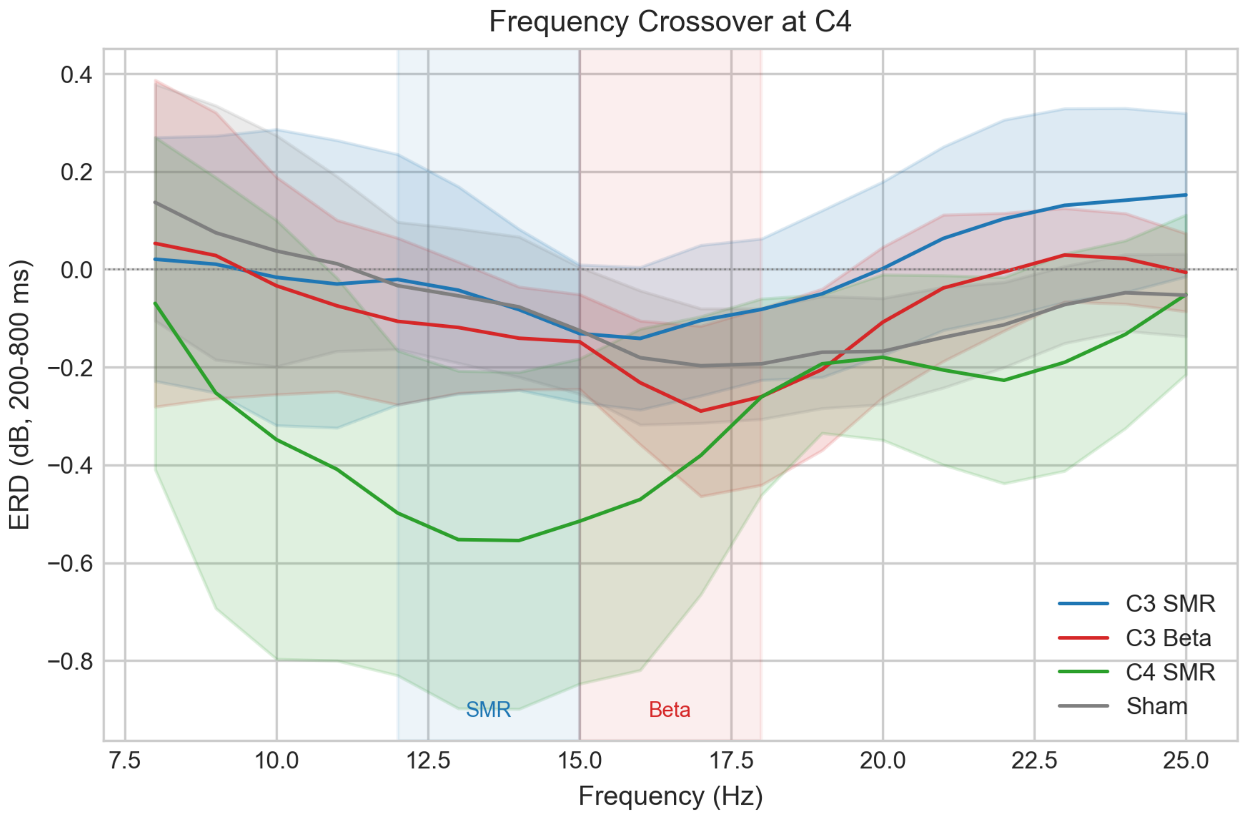


Figure 8: **Figure S8.** Frequency crossover at C4 (all 4 groups).

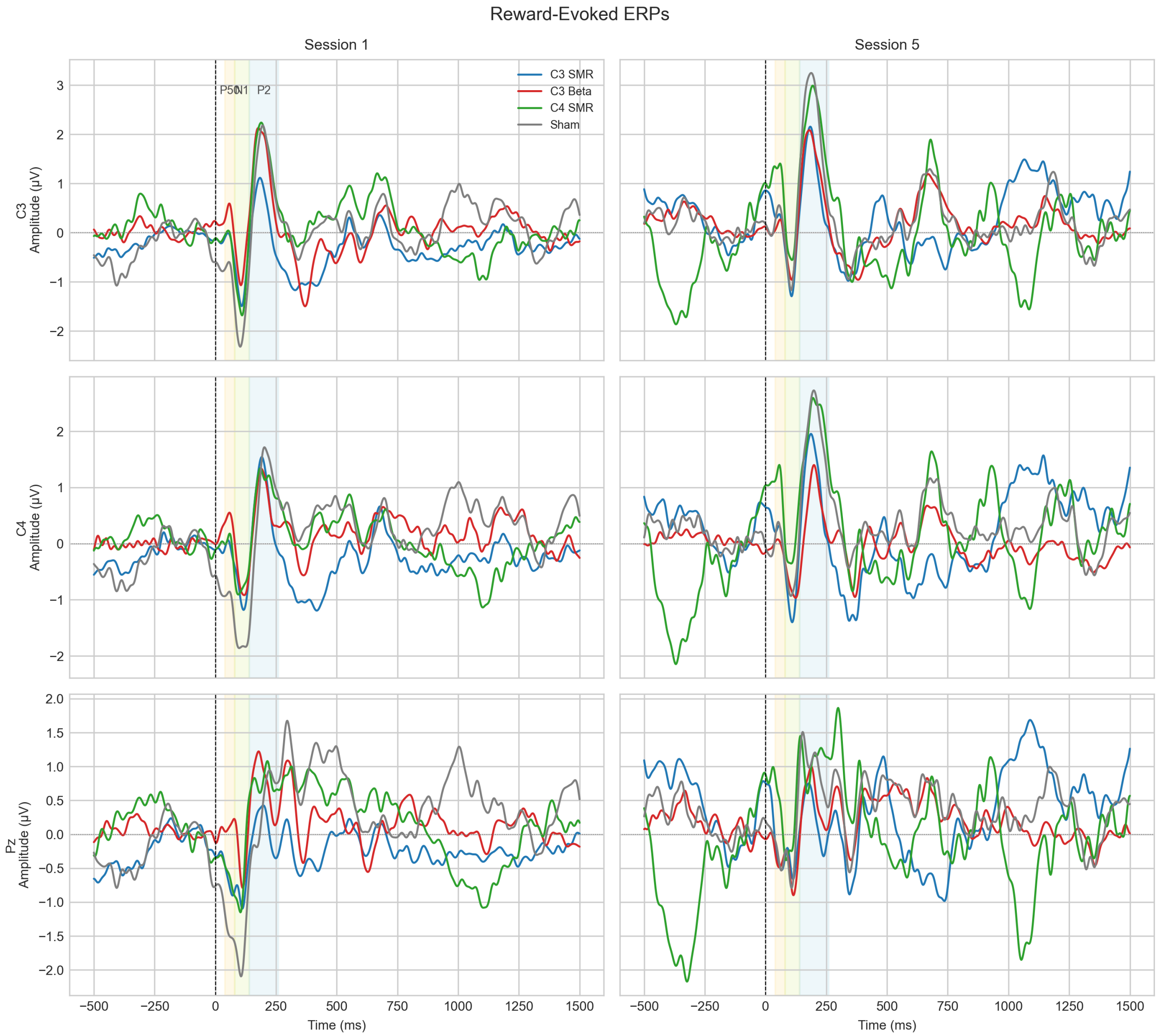


Figure 9: **Figure S9.** Full reward-evoked ERP waveforms at C3, C4, Pz (Sessions 1 and 5). Component windows shaded: P50 (40–80 ms), N1 (80–140 ms), P2 (140–260 ms).

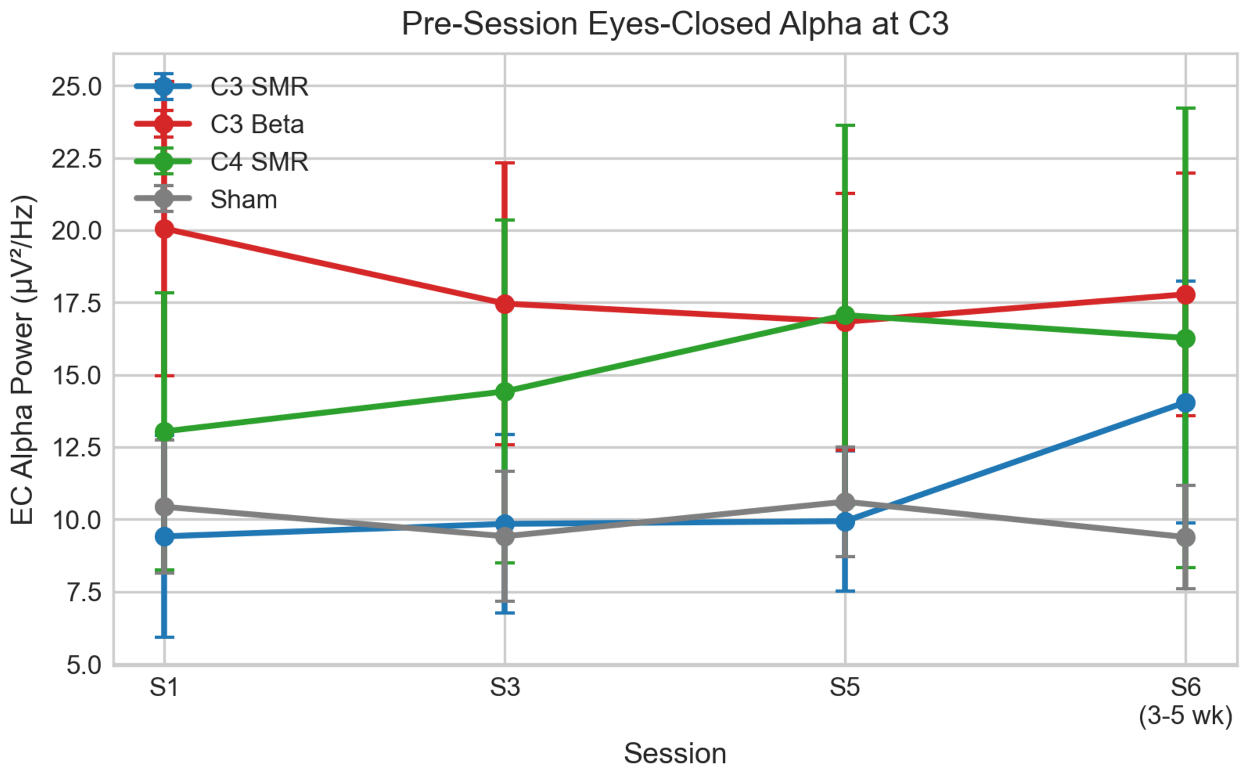


Figure 10: **Figure S10.** Pre-session eyes-closed alpha power at C3 across sessions. SMR groups show cumulative accumulation; sham and C3 Beta are flat or declining. Error bars: SEM.

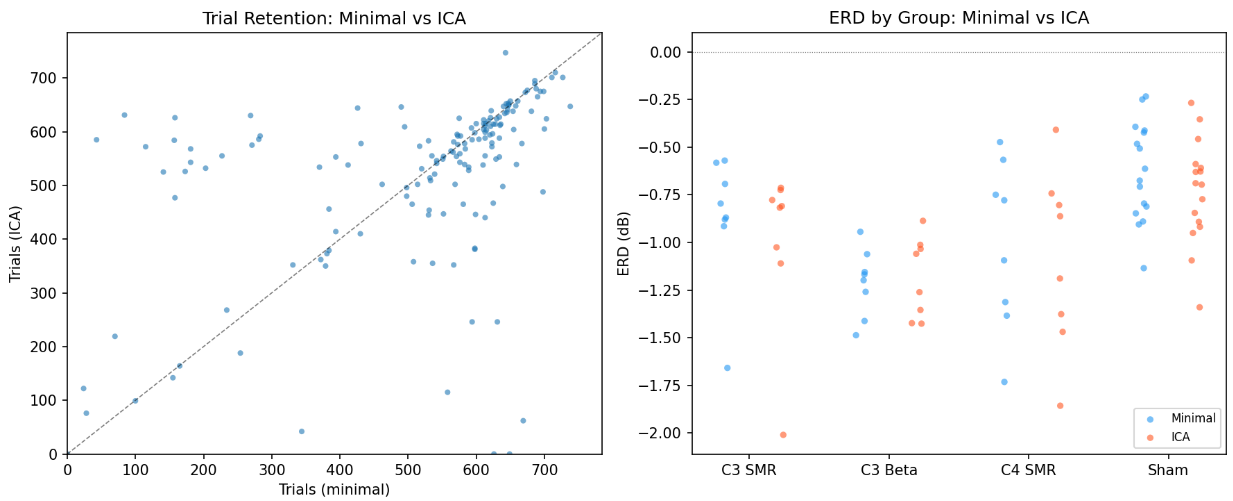


Figure 11: **Figure S11.** Sensitivity analysis: minimal vs ICA preprocessing. Left: trial retention scatter plot (each point = one subject × session); the dashed line indicates equal retention. Right: per-group ERD distributions under both pipelines. The minimal pipeline produces slightly larger effect sizes despite comparable trial counts, consistent with signal preservation (Delorme, 2023).

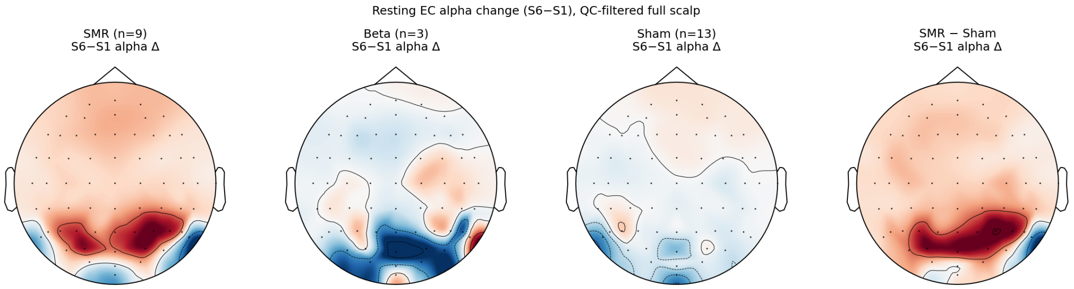


Figure 12: **Figure S12.** Exploratory 64-channel topography of resting eyes-closed alpha change (S6 minus S1), QC-filtered (25 of 38 recordings retained). Four topographic maps show the group-mean S6-minus-S1 alpha power change for SMR ($n = 9$), Beta ($n = 3$), Sham ($n = 13$), and the SMR-minus-Sham difference. The SMR group shows a broad posterior increase; the difference map concentrates the effect over right parietal-occipital scalp sites. The near-significant cluster ($p = 0.082$; Pz, C2, C4, CP2, CP4, P2, P4) does not survive correction. This analysis is exploratory and hypothesis-generating only.
